# Do macroinvertebrate abundance and community structure depend on the quality of ponds located in peri-urban areas?

**DOI:** 10.1101/2023.10.20.563281

**Authors:** Florence D. Hulot, Christophe Hanot, Sylvie Nélieu, Isabelle Lamy, Sara Karolak, Ghislaine Delarue, Emmanuelle Baudry

## Abstract

Contamination is one of the major threats to freshwater biodiversity. Compared to other aquatic ecosystems, peri-urban ponds are unique because they are embedded in human-dominated areas. However, it is poorly understood how different land uses such as urban or agricultural contribute multiple pollutants to ponds and thus affect pond biodiversity. In this work, 12 ponds located in a peri-urban area (Ile-de-France region, France) were monitored for 2 consecutive years in spring and fall. We surveyed macroinvertebrates and measured the physicochemical parameters and contaminants of different classes (trace elements, pharmaceuticals, pesticides and polycyclic aromatic hydrocarbons) in both water and sediment. The objective was twofold: (1) to explore local and regional macroinvertebrate spatiotemporal diversity and (2) to understand the effects of contaminants on community structure. We observed 236 macroinvertebrate morphotaxa, none of which were rare or sensitive to pollutants. Morphotaxa richness showed small differences between ponds but no difference between there was no effect of field campaign. There was no effect of ponds and field campaign on morphotaxa diversity and equitability. We did not observe a relationship between land use around the pond (agricultural, urban, or semi-natural) and diversity indices with the exception of the proportion of agricultural land in the vicinity of the pond on equitability. Regional beta diversity (between ponds) showed that differences in morphotaxa composition reflected species replacement more than differences in species richness; these were primarily due to the high abundance of pollutant-tolerant species in some of the ponds. The effects of environmental parameters on community structure were studied using partial redundancy analysis based on the presence-absence of morphotaxa, showing that community assemblages are shaped by sediment levels of pharmaceuticals, water conductivity and ammonium concentration. In conclusion, ponds in peri-urban areas are exposed to various human activities, with our results suggesting that this exposure leads to chronic and diverse contaminations that affect morphotaxa communities.

## 1. Introduction

Freshwater ecosystems, including lakes, reservoirs and rivers, host around 9.5% of all described species despite representing approximately 2.3% of the world’s surface area and only 0.01% of its water (Reid et al. 2019). They therefore deserve special attention in terms of their protection and conservation. In a recent review, Reid et al. (2019) listed 12 emerging or intensifying threats to freshwater biodiversity. Among the different freshwater ecosystems, ponds are particularly vulnerable because of their small size (Biggs et al. 2017), although they are understudied probably on account of their presumed insignificance (Biggs et al. 2017; Cereghino et al. 2008). Several studies showed that small water bodies such as ponds, ditches and streams host a higher biodiversity than large water bodies (Biggs et al. 2017). In particular, because of their isolation or small size, they may have low local alpha diversity. However, a high dissimilarity between these aquatic systems may lead to high regional beta diversity (Clarke et al. 2008; Davies et al. 2008; Williams et al. 2004). The small habitat size of ponds may also support rare and scarce species. Scheffer et al. (2006) showed that shallow lakes and ponds support a higher richness of aquatic birds, plants, amphibians and invertebrates than large water bodies. This higher richness is accompanied by a low diversity of fish, if not their complete absence (Scheffer et al. 2006).

Peri-urban areas are characterised by complex landscapes with both agricultural and urban covers and a mixture of different uses and users (Poggi et al. 2021; Zoomers et al. 2017). They are typical zones of continuous transformation (Zoomers et al. 2017) or “restless landscape” (Friedmann 2016). However, ponds and wetlands in peri-urban environments are understudied (Wanek et al. 2021). Although urban ponds are less diverse than rural ponds, they may host threatened species, thus advocating for good management practices (Oertli and Parris 2019). Likewise, motorway stormwater retention ponds can play a significant role in macroinvertebrate diversity at the regional level (Le Viol et al. 2009; Meland et al. 2020).

Contamination by pollutants is one of the 12 threats to freshwater biodiversity identified by Reid et al. (2019). Ponds are subjected to multiple contaminations depending on their environment, whether in an agricultural, urban or highway setting. These stressors have been studied both independently and in combination and include, among others, pesticides (Trigal et al. 2007), major ions, nutrients and wastewater-associated micropollutants (Berger et al. 2018), polycyclic aromatic hydrocarbons (PAH) (Uher et al. 2016), pharmaceuticals and heavy metals (Andreu et al. 2016), metals, ions and PAH in combination (Sun et al. 2018). The distinctive feature of peri-urban ponds is that they are embedded in a human-dominated matrix with different activities, which can be the source of multiple stressors. In peri-urban areas, although ponds are not located far from each other, they might be exposed to different dominant stressors or combine different categories of pollutants because of their small catchment area (Biggs et al. 2017; Cereghino et al. 2008). In addition, contaminants in ponds may change with time if the surrounding landscape is submitted to changes such as urbanisation or temporary construction sites, or if it is exposed to new materials such as nanomaterials or personal care product additives. Ponds may accumulate these contaminants, particularly in their sediments and, consequently, freshwater life is affected by changing contaminant cocktails.

Aquatic macroinvertebrates encompass a rich and diversified set of taxa that are universally found in freshwater ecosystems. They exhibit a wide range of sensitivity to environmental stressors and, as a consequence, their local diversity and abundance are commonly used as indicators of perturbations (Sumudumali and Jayawardana 2021; Tachet et al. 2010). Aquatic macroinvertebrates are mostly sedentary, at least during their larvae stage. They inhabit different habitats and their life cycle, for the majority of macroinvertebrates, is annual (Tachet et al. 2010). For these reasons, these organisms are good indicators of pollution, as the recolonisation of perturbed areas takes time. Macroinvertebrate indices of water quality are based on the presence-absence or abundances of macroinvertebrates. However, in addition to the local diversity known as alpha diversity, spatial and temporal diversity give information respectively about the spatial structuration and temporal changes of communities (Legendre and Condit 2019). Thus, their spatial and temporal beta diversity might prove useful to assess the impacts of stressors in changing peri-urban areas.

Here we report the results of a study in which we monitored 12 ponds in a peri-urban area located in the Ile-de-France region (France). The chosen ponds were characterised by different proportions of agricultural, urbanised, grassland and forest surfaces in a 100-m radius buffer. We therefore aimed to link land use, contaminant concentrations in water and sediment as well as macroinvertebrate distribution in ponds. Our objective was to understand whether water and sediment pollutants and land-use characteristics are constraints to macroinvertebrate distribution in ponds. To do so, we sampled macroinvertebrates and measured pond quality parameters, including various urban and agricultural contaminants, to quantify the main pesticides, pharmaceuticals, PAH and trace elements (TE) in water and sediments. Depending on their chemical properties, contaminants may accumulate either in water or in sediment. We therefore monitored both compartments. Nélieu et al. (2020) showed that the water contamination profiles of these ponds differed depending on their location, and that the agricultural landscape explained these differences more than urban land uses. In some of the ponds, the environmental risk exceeded the thresholds of risk quotient mainly due to pesticides (Nélieu et al. 2020). Here, we analysed the macroinvertebrate communities and hypothesised that (1) local macroinvertebrate diversity is higher in ponds located in environments dominated by grasslands and forests than in those dominated by agriculture and urban areas; (2) ponds hosting rare and pollutant-sensitive macroinvertebrate morphotaxa highly contribute to regional diversity; and (3) water and sediment contaminants influence morphotaxa distribution in ponds.

## 2. Material and methods

### 2.1. Study area

The selected study area is the Saclay Plateau (N: 48° 43’ 59.99" E: 2° 10’ 0.01), located in the junction zone between the Parisian agglomeration and its large surrounding plains. Until recently, the territory had a mainly agricultural vocation, although the ongoing development of a scientific and technological pole in the area has considerably increased the urban hold on the territory. At the same time, the Natural, Agricultural and Forest Protection Zone of the Saclay Plateau extending over more than 4,000 ha was created in 2010, thus perpetuating the agricultural use of land on the plateau. The Saclay Plateau thus presents major challenges in terms of the coexistence of urban and agricultural areas and biodiversity in a context of growing urbanisation. Within this plateau, we selected 12 ponds with surface areas ranging from 64 to 828 m^2^ (mean ± SD: 540 ± 320 m^2^; median: 566m²) and with different potential exposures to agricultural and urban activities. The proportion of agricultural and urbanised surface around each pond was calculated in a 100 m radius buffer. As the aim was to study the effect on water quality of land use in the vicinity of ponds in a fairly fragmented landscape, we used a short-radius buffer zone. To calculate the different types of surface, we used the map of mainland France land cover produced by the Theia land data services using Sentinel 2 and Landsat 8 data (Theia 2017). The proportion of urbanised land was calculated by adding together three Theia categories: dense built-up areas, diffuse built-up areas and industrial and commercial areas. The proportion of agricultural land was calculated by adding two Theia categories: winter crops and summer crops. The remaining areas consist of forest and grassland. In the following, we refer arbitrarily to ponds A to L.

### 2.2 Sampling methods for invertebrates and for water and sediment

The sampling of invertebrates as well as water and sediment was systematically performed at the same points in the 12 ponds. To ensure that water and sediment sampling and invertebrate collection did not interfere with each other, they were carried out with a time lag of about 1 week.

A preliminary analysis was conducted to determine the most appropriate timing for sampling the invertebrates in the ponds. Several ponds were sampled every month from May to October 2015 inclusive, which are the periods during which most species are active in our geographical area. Eight of these ponds were included among the 12 ponds selected in our study. Non-metric multi-dimensional scaling (NMDS) analysis of the collected taxa showed that sampling in June and September covered almost all the diversity of the macroinvertebrate species present in the ponds (Hanot, unpublished results). Sampling was thus performed in June and September during 2016 and 2017 in the 12 ponds selected for the analysis. In the following, we refer to the field campaigns as C1 (2016 June), C2 (2016 September), C3 (2017 June) and C4 (2017 September).

Macroinvertebrate sampling was carried out on two different main habitats on either side of each pond to best represent the biodiversity of the ponds in a standardised manner. Sampling was made using a pond-net with a 77 cm fiberglass handle, a trapezoid frame measuring 39 x 14 cm x 32 cm, a 1 x 1 mm mesh and a pocket depth of 40 cm. Samples were collected by making an infinity symbol by hand eight times (i.e., a figure eight on its side), ending with a fast move from the bottom to the top in the axis point of the symbol, to collect organisms trapped in the vortex created by the sequence of movements. During this sequence, the frame of the net pond was a few centimetres above the substratum, which allowed us to lift and collect the benthos thanks to the upward current created by the “infinity” movement. To sample each pond in the same way, the same operator made the movements at an arm’s distance from the bank. In a small number of cases, the water level in the ponds was too low to sample from the selected habitat. In this case, no collection was made at this point or at any other to ensure that the samples were collected in the same location during the entire study period.

During the collections in 2016, the content of the net pocket was quickly placed in a white basin measuring 80 x 40 x 10 cm. All specimens were sorted by eye and fixed in absolute ethanol 99% (Fisher Chemical, CAS 64-17-5) using entomological forceps. Collection stopped when 5 min had elapsed without seeing any moving specimen. Thereafter, all samples were kept at 7°C until the identification stage. During the 2017 campaigns, to speed up the sample collection, water was squeezed from the net pocket as gently as possible to avoid destroying the specimens, and the result was then fixed in absolute ethanol 99% in suitable containers and kept at 7°C. Specimens were sorted using a Motic SMZ 171 binocular microscope and entomological forceps, fixed again in absolute ethanol 99% and then kept at 7°C. Our sampling protocol resulted in 88 invertebrate samples: 12 ponds x 2 points x 2 years x 2 seasons, minus 8 cases (4 ponds x 2 points) when the collection was not possible due to a low water level. Invertebrate identifications (see next Section) were carried out separately for each sample.

In 2016 and 2017, in each of the 12 ponds, water and sediment samples were collected in spring and autumn from the same two points selected for invertebrate sampling. The water samples were taken approximately 1 m from the edge of the ponds using a stainless steel beaker with an extendable handle. Sediment sampling was performed with the same device used to collect the water samples.

### 2.3 Identification of invertebrates

Different books (Bameul 1985; Guignot 1947; Hansen 1987; Holmen 1997; Jansson 1986; Olmi 1976; Poisson 1957 ; Tachet et al. 2010) and websites (http://www.perla.developpement-durable.gouv.fr/, http://coleonet.de) were used as a reference to perform the morphological identification of all specimens under a Motic SMZ 171 stereo microscope. Identifications were made at the lowest possible taxonomic level. Specimens that could not be identified at the species level were identified at a higher taxonomic level, while adding a numerical suffix when more than one species was present (e.g., *Microvelia* sp.2). Because of these different levels of determination, we hereafter refer to specimens as “morphotaxa”, which are defined as taxa that share the same morphological characteristics. When it was not possible to link the different stages (larva, nymph, adult) to the same species, they were assigned to different morphotaxa. Some specimens of each taxon were kept in tubes of 2 mL, 5 mL or 40 mL according to their size, in absolute ethanol 99%, to be used as a reference. This made it possible to constitute a reference base for the invertebrates in the ponds by linking the reference specimens with their morphotaxon names. Each specimen of each sample was then identified using books, websites and the reference base. All specimens were counted by morphotaxa for each sample.

### 2.4 Determination of water and sediment quality parameters, including trace elements and organic pollutants

The choice of contaminants assessed here was based on the local activities: cereals, maize, rapeseed, sunflower, orchard and vegetable crops for pesticides; nearby roads for TE and PAH; and the presence of humans, farms, and domestic pets for pharmaceuticals. Samples were used to determine the main physicochemical parameters, including major and TE as well as organic contaminants (PAH, pesticides and pharmaceuticals) as described in Nélieu et al. (2020).

For the water samples, the following measurements were taken directly on site with probes: pH, conductivity, temperature, dissolved oxygen (DO) and turbidity. Other data were obtained rapidly in the laboratory (mainly within one day of sampling) using standardised methods: dissolved organic carbon (DOC) by thermic oxidation and IR analysis of carbon dioxide, chemical oxygen demand (COD, norm NF EN ISO 15705), suspended solids (SS, norm NF EN 872), nitrates (NO ^-^, norm NF EN ISO 10304-1), nitrites (NO ^-,^, NF EN ISO 26777), total nitrogen (TN) from the addition of Kjeldahl nitrogen (Kjeldahl method, norms NF EN 25663 and NF EN ISO 11732) with nitrates and nitrites, total phosphorus (P, norm NF EN ISO 15681-2 and NF EN ISO 6878.), anions (norm NF EN ISO 10304-1) and cations (norm NF EN ISO 14911), as well as major and TE (norm NF EN ISO 17294-2: Al, As, B, Be, Cd, Cr, Cu, Fe, Hg, Mn, Ni, Pb, Sn, U and Zn), 15 PAH, pesticides (25 herbicides, 1 safener, 7 fungicides and 2 insecticides) and 12 pharmaceuticals compatible with the multi-residue method applied after sample conservation at -20°C (see the exhaustive list in Table S1). Details on the methods used for this determination can be found in Nélieu et al. (2020).

Sediment samples were used to determine the contents in organic carbon (Corg), total nitrogen (N) and thus C/N ratio (norms ISO 10694 and ISO 13878) as well as total major and TE (after HF mineralisation and then ICP-AES or ICP-MS analysis according to the norm NF X 31-147/NF ISO 22036 - 17294-2), for the following: Cr, Cu, Ni, Zn, Co, Pb, Cd, Tl, Mo, Al, Ca, Fe, K, Mg, Mn, Na, P (P_2_O_5_), Bi, In, Sb and Sn. All measurements were made in the Laboratory of Soil Analysis of INRAe (Arras, France). The polycyclic aromatic hydrocarbons were analysed according to the European standard NF EN 16181 (2018) by pressurised liquid extraction and HPLC-fluorescence quantification. The same pesticides, metabolites and pharmaceuticals selected for analysis in water (Nélieu et al., 2020) were also monitored in sediments (see Appendix 1 for the analysis methods used for sediments).

Not all the measured pesticides and pharmaceuticals were detected in water and sediment (no detection or values below the quantification limits). Therefore, our study is based on fifteen herbicides (Atrazine, Atrazine-desethyl, Simazine, Terbuthylazine, Terbuthylazine-desethyl, Clomazone, Diflufenican, Napropamid, Acetochlore, Alachlore, Dimethachlore, Metolachlore, Chlorsulfuron, Metsulfuron-methyl and Nicosulfuron), seven fungicides (Boscalid, Dimoxystrobine, Epoxiconazole, Hexaconazole, Metconazole, Picoxystrobine, Tebuconazole), two insecticides (Imidacloprid and Pyrimicarb) and one pharmaceutical (Carbamazepine).

All the data were added to the In.Do.Res repository (https://doi.org/10.48579/PRO/TX0PU9,).

### 2.5 Statistical analysis

Prior to the data analysis, the invertebrate samples collected from both points in each pond were pooled. This allowed us to produce a contingency table containing the number of each morphotaxon for each pond and each sampling date. For the water quality parameters, TE and organic pollutants, we computed the mean of the two sample point values. Although the surface area of ponds is important, as pointed out in the introduction, it could not be included as a factor in the analyses because we only have a single measurement. All statistical analyses were performed with R 3.6.1 (Team 2020).

We performed analyses based on morphotaxa abundance and presence-absence. However, due to differences in specimen determination levels, the analyses may be biased and should therefore be taken with caution. To explore invertebrate community diversity, we computed the morphotaxa richness (alpha diversity), Shannon diversity index and Pielou evenness for each pond and each field campaign with the specnumber function in vegan 2.5-6 (Oksanen et al. 2019). We tested the effect of individual ponds and field campaign on these parameters with an analysis of variance with additive effects of pond and field session followed by a pairwise comparison with Tukey’s HSD test.

To study the dissimilarities between invertebrate communities, we computed the total beta diversity within each field campaign across all ponds using the beta.multi function of the betapart package 1.5.2 (Baselga et al. 2020). This function partitions the total beta diversity into two additive components, turnover and nestedness, which reflect species replacement and species richness difference, respectively (Baselga et al. 2020).

We calculated the contribution of each pond to beta diversity, that is, the local contribution to beta diversity (hereafter LCBD) following Legendre and Cáceres (2013) using the beta.div function of the adespatial package 0.3-8 (Dray et al. 2019). As pond dissimilarity may be different if computed with the abundance or presence-absence (PA) of morphotaxa, we calculated both using the Hellinger and Jaccard dissimilarity coefficients, respectively. The LCBD values, which represent the uniqueness of a pond in terms of taxa composition, were tested for significance with the null hypothesis of a random distribution of species among ponds within a sampling campaign (Legendre and De Cáceres 2013). We also computed the species contributions to beta diversity (hereafter SCBD) to identify the morphotaxa that contribute the most to beta diversity. This last index is calculated only with morphotaxa abundance. We explored the temporal effects (year and season) on morphotaxa assemblages but as the results are not robust, we present them in Appendix 2.

To explore the relationship between the parameters describing diversity and land use, we tested the effects of the proportion of urban, forested and agricultural areas on morphotaxa richness, diversity, equitability and LCBD with a linear mixed effect model with the field campaign as a random effect. We used the lme4 package, v.1.1-23 (Capps et al. 2015).

Finally, we explored the relationship between the environmental parameters and the macroinvertebrate assemblages to identify the parameters that best explain the community structures. The concentrations of water TE, PAH and pharmaceuticals were very low, being at the detection limit; for this reason, we did not include them in the analyses. Prior to the analyses, to reduce the high number of TE in the sediments, we performed principal component analysis (PCA) in all ponds for the four field campaigns. The first two axes of the PCA explain 68.67% of the variability (Figure S1). The ponds are arranged on the first axis (TE1) from low to high TE concentrations. The second axis (TE2) discriminates ponds with high concentrations of TE (Sb, Cd) from those with high concentrations of major elements (Na, Mg, Fe). We collected the coordinates of each pond-field campaign combination for the first two axes of the PCA and then used them in the following statistical analyses as summaries of each contaminant group effects on ponds. We summed the concentration of the different PAH and used the total PAH concentration in the following analyses. We used redundancy analysis (RDA) (Borcard et al. 2011) and ran two analyses, one with the Hellinger-transformed abundances of morphotaxa and another with the PA of morphotaxa. In the RDA, the response matrix is the abundance or PA of morphotaxa in all ponds and the four field campaigns, whereas the explanatory matrix is the environmental parameters, including the contaminants for the same pond-field campaign combination. We used the following parameters in the RDA: (1) for water: conductivity, suspended matter, COD, DOC, concentrations of TN and phosphorus, orthophosphate, ammonium, organic carbon, herbicides and insecticides; and (2) for sediments: herbicides, fungicides, insecticides, pharmaceuticals, PAH and the first two PCA axes performed with TE.

We used the rda function of the vegan package v2.5-6 (Oksanen et al. 2019). We tested the significance of the two RDA results by the permutation of the overall analysis and each axis. The two RDA were significant with a threshold level of 5%. We tested for linear dependencies among the explanatory variables and computed the variance inflation factors (VIF) of the variables with the vif.cca function. We computed the adjusted R² with the RsquareAdj function. The VIF was very high for several explanatory variables and so to reduce the correlations between them, we computed a forward selection of the explanatory variables using the forward.sel function. The method produces parsimonious models, which we have tested by permutation and VIF.

## 3. Results

### 3.1 Morphotaxa richness, Shannon diversity index and evenness

In total, we identified 236 morphotaxa, which represent a total of 22 orders and 54 families including 13 orders and 42 families for arthropods and 8 orders and 37 families for insects (some morphotaxa could not be assigned to an order or family). The Baetidae *C. dipterum* was ubiquitous during campaigns C1, C2 and C3 but totally absent in C4. Morphotaxa richness ranged from 7 to 49 with a median value of 25. Statistical analyses show a weak effect of pond on morphotaxa richness (Figure 1 and Table S2). Morphotaxa richness was significantly higher in pond J than in ponds E (p=0.023) and I (p=0.035). Morphotaxa richness changed between field campaigns (p≤ 0.001, Table S1), being significantly higher in C1 and C2 than in C4 (respectively p=0.004 and p=0.008). The Shannon index and evenness ranged from 1.07 to 2.9 and from 0.33 to 0.82, respectively. The field campaigns and ponds had no effect on the Shannon index. Only the field campaign had an effect on evenness (p=0.02) with a higher evenness in C4 than in C1 (p=0.02).

**Figure 1.**
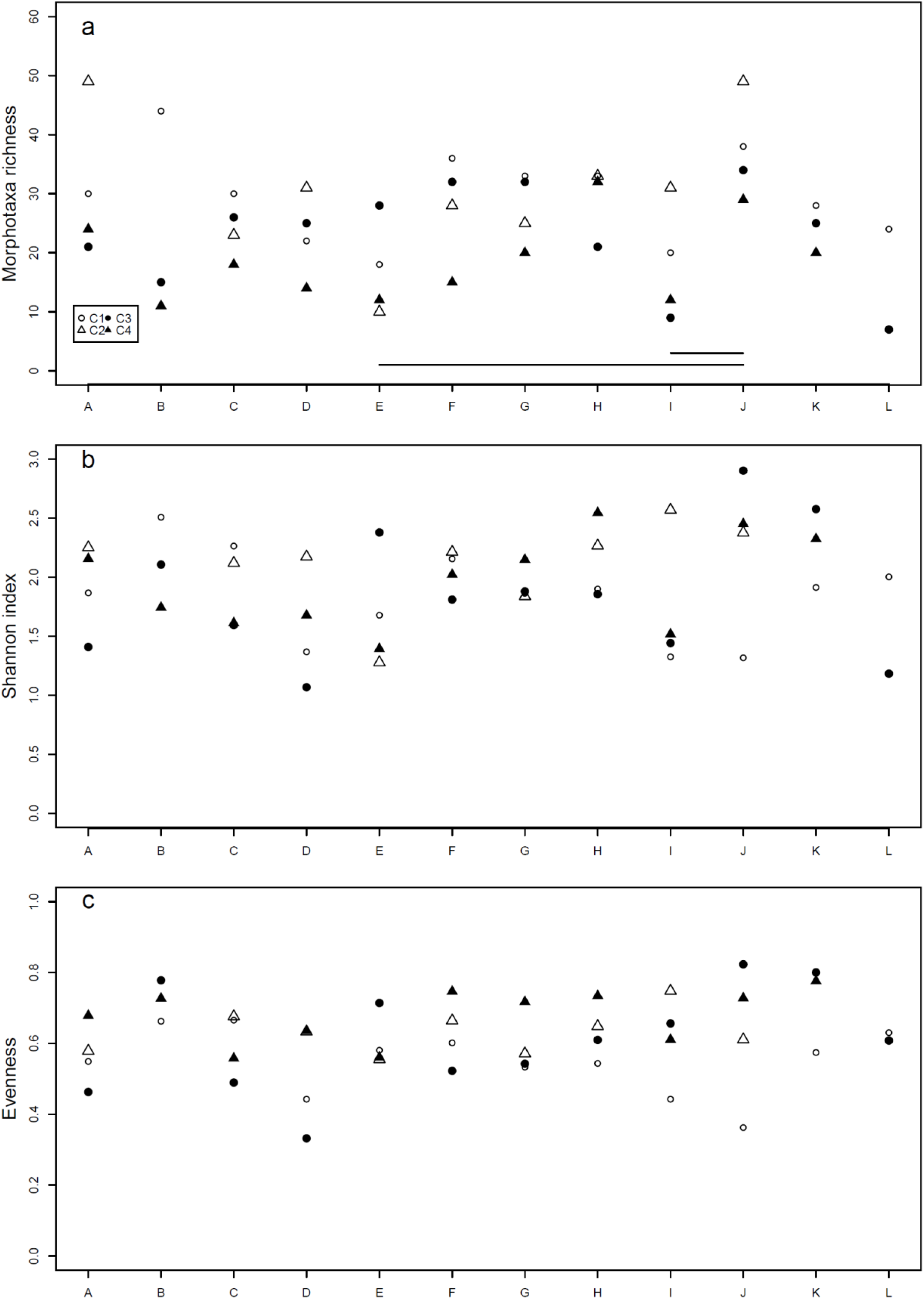
Morphotaxa richness (a), Shannon index (b) and evenness (c) in the 12 ponds for the four field sessions. The legend for the four field sessions is given in panel (a). The horizontal segments in panel (a) link the ponds significantly different.

### 3.2 Beta diversity: Spatial dissimilarities between ponds

The total beta diversity based on morphotaxa PA was very similar for each field campaign with values between 0.92 and 0.93 (Table S3; for comparison, results for analyses based on morphotaxa abundances are in Table S2). The turnover, which reflects the level of species replacement between ponds as opposed to species loss, represents between 88% and 90% of this total beta diversity. The LCBD of each pond based on morphotaxa PA varied across the field campaigns (Figure 2, Table S3). Pond E has a significant contribution to regional biodiversity in C1 and C2. Pond I makes a significant to highly significant contribution during the four field sessions and ponds J and L in C4 and C3, respectively.

**Figure 2.**
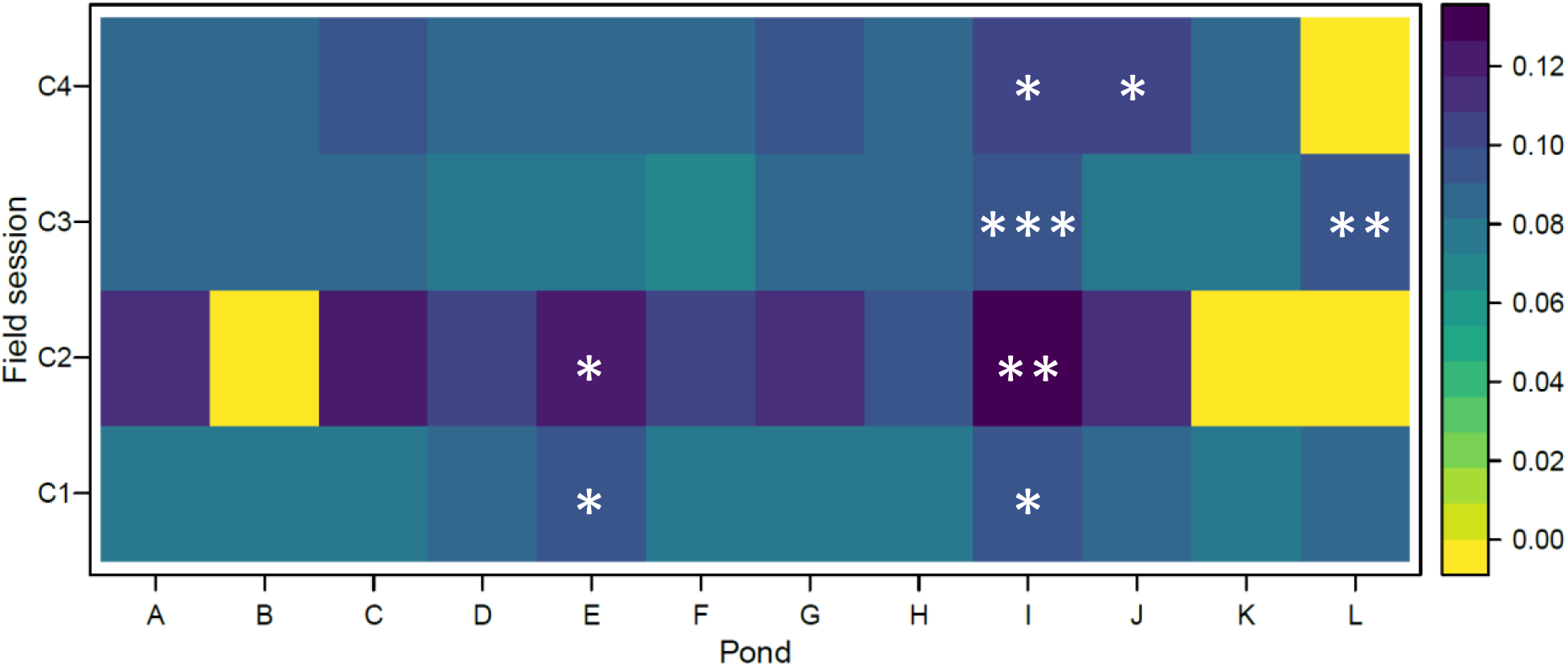
Local contribution to beta diversity (LCBD) of each pond for the four field sessions. The LCBD is computed with the presence-absence of morphotaxa. The LCBD are computed independently for each field session. The symbols show the significant contributions with: ***: p ≤ 0.001; **: 0.001<p ≤ 0.01; *: 0.01 < p ≤ 0.05.

To shed light on the LCBD results, we briefly present the Species (here morphotaxa) Contribution to Local Biodiversity (SCBD) based on morphotaxa abundances. The analysis reveals that a relatively low number of morphotaxa explains most of the dissimilarities among ponds. The 15 morphotaxa with the highest contributions are listed in Table S5 and the comprehensive list of morphotaxa SCBD is provided in Table S6. The 15 morphotaxa with the highest contributions account for 83%, 73%, 71% and 70% of the total SCBD in the C1, C2, C3 and C4 field campaigns, respectively. Some are common to all four campaigns (*Asellus* sp., two different *Chaoborus* sp., different Chironomini morphotaxa) or to three campaigns (*Cloeon dipterum*; Clitellata; *Physella acuta*; *Valvata macrostoma*).

### 3.3 Relationship between land use, environmental parameters and macroinvertebrate assemblages

The linear mixed effect model testing the effects of the different types of land use around the ponds on parameters characterising the macroinvertebrate diversity showed that equitability increased significantly with the proportion of agricultural area in pond vicinity (p=0.026). Morphotaxa richness and diversity, and LCBD were not significantly affected by land use.

To clarify the relationship between the environmental parameters for water and sediment and the macroinvertebrate assemblages, we ran redundancy analysis (RDA) with morphotaxa PA. The initial model with all the environmental parameters was significant (p = 0.009). As some of these parameters had strong collinearities (as shown by the high VIF values), a forward model selection procedure was used to obtain a more parsimonious model. The final model describing the morphotaxa PA contained three explanatory variables: conductivity, pharmaceutical concentration in sediments and ammonium concentration in water (Table 1). The parsimonious model is highly significant (Df = 3, 40, F-value = 1.740, p=0.002) with no strong collinearity between the variables (all VIF are around 1) and is reduced to one significant canonical axis (Figure 3). Pond C is associated with high concentration of pharmaceuticals in the sediment; pond G is also associated with high concentration of pharmaceuticals in the sediment in addition to high water conductivity.

**Figure 3.**
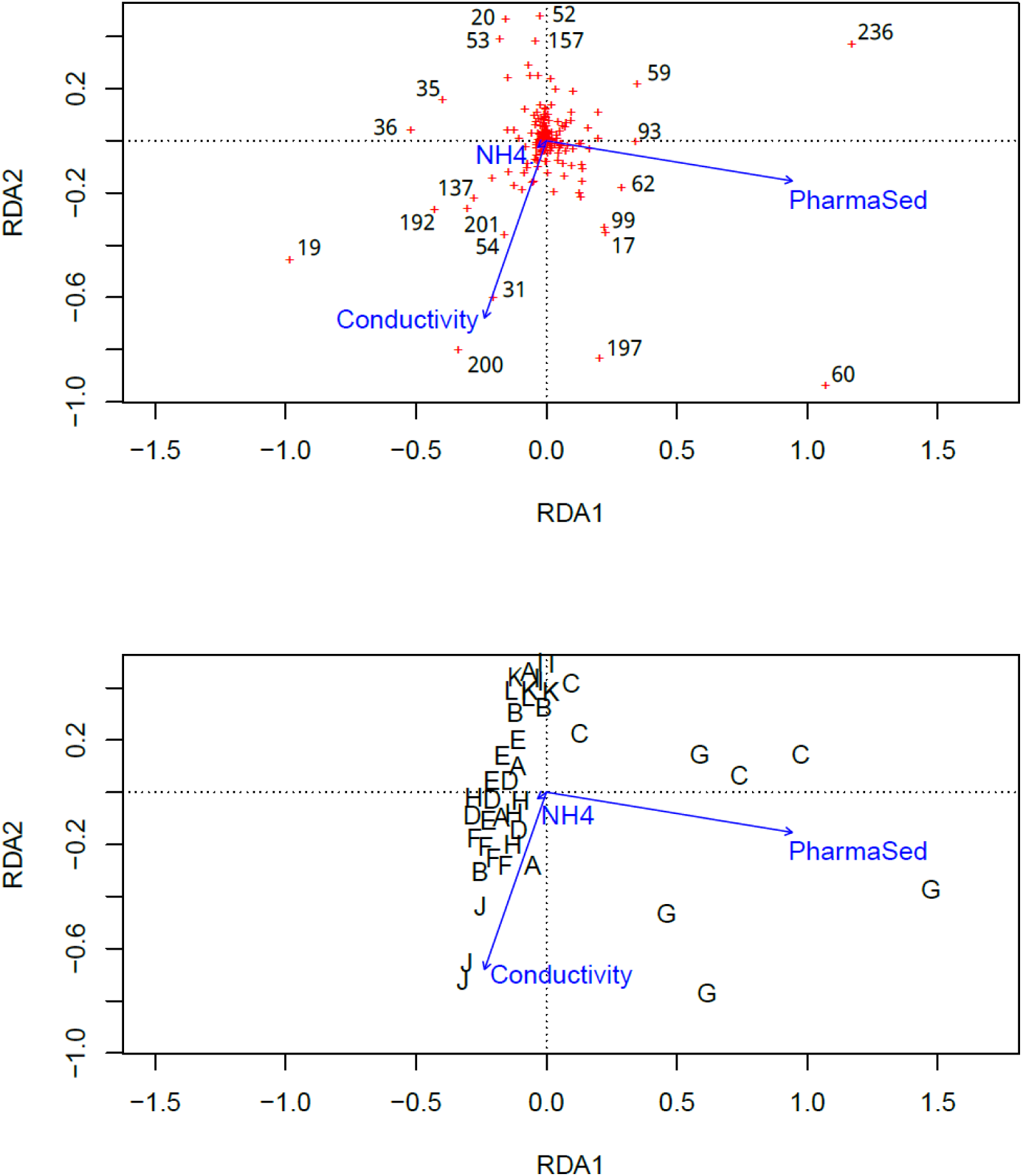
Parsimonious redundancy analysis (RDA) based on morphotaxa presence-absence. The biplots show the variables (blue arrows) with either the morphotaxa (numbers, panel a) or the ponds (letters, panel b). The number of morphotaxa is shown in brackets: Anophelinae 01 (17); *Asellus* sp. (19); Baetidae 02 (20); Ceratopogoninae 02 (31); *Chaoborus* sp. 01 (35); *Chaoborus* sp. 02 (36); Chironomini 04 (52); *Chironomus* sp. 01 (53); *Chironomus* sp. 02 (54); Clitellata 01 (59); *Cloeon dipterum* (60); *Coenagrion* sp.01 (62); *Dugesia* sp. 2 (93); *Erythromma viridulum* (99); Hesperocorixa 03 (137); *Hygrotus inaequalis* (157); *Physella acuta* (192); *Plea minutissima* (197); *Potamopyrgus antipodarum* (200); *Proasellus* sp. (201); and *Valvata macrostoma* (236). Conductivity: water conductivity; PharmaSed: concentration of pharmaceuticals in sediment; NH4: water ammonium concentration.

**Table 1.** Results of the redundancy analyses (RA) parsimonious models.

| Variables | Cumulative adjusted R<sup>2</sup> | F value | *p* value |
| --- | --- | --- | --- |
| Water conductivity | 0.032 | 2.401 | 0.001 |
| Sediment pharmaceuticals | 0.057 | 2.111 | 0.001 |
| Water ammonium concentration | 0.077 | 1.922 | 0.006 |

Pond J is associated with high water conductivity. Ponds, A, B, D, E, F, H, I, K and L are less affected by environmental parameters although ponds K and L seem associated with low water conductivity. A few morphotaxa stand out and are associated with some environmental variables (morphotaxa scores in the RDA are in Table S7). *Valvata macrostoma* is associated with high pharmaceutical concentrations in sediment; *Potamopyrgus antipodarum* is associated to high water conductivity; *Cloeon dipterum* and *Plea minutissima* are associated to high values both variables. Baetidae 02, *Chironomus* sp. 01, *Chironomus* sp. 02 and *Hygrotus inaequalis* are associated with ponds in which these environmental parameters have low values.

## 4. Discussion

### 4.1 Morphotaxa distribution in the ponds and effects of land use

We changed our experimental design for macroinvertebrate sampling between the 2 years of the study. The results show a greater morphotaxa richness in the first year compared with autumn in the second year. The morphotaxa diversity was not affected and the eveness slightly affected by the field campaign. As a consequence, it is difficult to conclude on the effects of the protocol change, as the effects may be small or may have been buffered by an annual effect.

Our analysis is based on morphotaxa determined at different taxonomic levels, which is open to criticism. The aim of our work is to compare the response of assemblages to the presence of pollutants and not to compare the diversity of the ponds studied with other ponds. In this sense, questions of determination level are less important, since the same precision has been maintained for all samples. Furthermore, studies on interaction networks have shown that the level of determination of specimens has little effect on network characteristics, provided that this level of determination does not fall below too high a threshold (Llopis-Belenguer et al. 2023; Renaud et al. 2020).

Among the 236 macroinvertebrate morphotaxa identified, we did not find endangered, vulnerable or even rare species. However, we observed exotic species: the molluscs *Potamopyrgus antipodarum* and *Physella acuta* and the crayfish *Procambarus clarkii* listed as an invasive alien species in the European Union (European Union 2016). Molluscs and crustaceans are the most frequent freshwater macroinvertebrate invaders (Oertli and Parris 2019; Patoka et al. 2017). The pet trade is one of the main introduction pathways, and both mollusc species can “hitchhike” on intended shipments (Patoka et al. 2017). One individual *P. clarkii* was found in pond F in C3. Pond F also hosted the two exotic molluscs, and pond J hosted *P. antipodarum*. We should stress here that pond F, though in a forested area, is located on the Paris-Saclay university campus with heavily frequented paths in close vicinity. The campus is open to the public, which may favour the dissemination of invasive species.

Overall, our results show that the ponds distributed along an urbanisation gradient are quite dissimilar, as beta diversity relies mostly on morphotaxa turnover with a comparable morphotaxa diversity. No pond stands out consistently across the four sampling campaigns in terms of the morphotaxa contribution to regional diversity except for pond I and, to a lesser extent, pond E. The LCBD indicates the uniqueness of communities either because they are rich and host typical taxa or because they are degraded with a limited number of common taxa (Legendre 2014; Legendre and De Cáceres 2013). Here, the uniqueness highlighted by the significant LCBD points to degraded ponds. For instance, pond I has a low diversity, and its most striking feature is the absence of Baetidae in C4, whereas the mean abundance in other ponds was 81.2 individuals (± 47.0 SD). Despite the restrictions described above, we have calculated the SCBD, based on morphotaxa abundances, because it supports the idea that some ponds stand out because they are degraded. The lists of morphotaxa contributing the most to regional diversity encompass common morphotaxa, some of which are characteristic of degraded communities such as Chironomidae, *Chaoborus* sp., *C. dipterum*, and so on. For instance, in C2, pond C was characterised by high abundances of *Chaoborus* sp. 01, *C. dipterum*, *Corixa* sp. and *V. macrostoma*, and pond E by *C. dipterum*, Orthocladiinae and Tanytarsini 01. Pond L in C3 had a low diversity and was dominated by *Notonecta* sp. 01. The uniqueness of a community as shown by a high LCBD may also indicate the presence of invasive species (Legendre 2014). However, the two ponds hosting exotic species as well as an invasive species had no significant LCBD values.

We initially hypothesised that local macroinvertebrate diversity is higher in ponds located in rural areas than in those located in agricultural or urban areas. This hypothesis is supported by different studies (Blicharska et al. 2017; Johnson et al. 2013; Noble and Hassall 2015; Thornhill et al. 2017). However, our results do not support this hypothesis, as we found only an effect of the proportion of agricultural area on morphotaxa evenness. Our results likewise do not support the second hypothesis regarding rare and pollutant-sensitive morphotaxa, as we do not observe any of them. On the contrary, we found invasive and exotic species. LCBD values could help to identify these ponds (Legendre 2014), although they only identified pond J, probably because of the very high density of *P. antipodarum*.

### 4.2 Effects of environmental parameters on macroinvertebrate assemblages

Our results showed that among the numerous environmental parameters and pollutants measured in water and sediment, very few are critical for macroinvertebrate assemblages in ponds. The concentration of pharmaceuticals in sediment and water conductivity are the most structuring parameters of macroinvertebrate assemblages. Although these results should be treated with caution, the analysis with morphotaxa abundances reveal other important parameters as the concentration of fungicides in sediment as well as MTE1, insecticides, organic carbon, and COD in water. Nélieu et al. (2020) highlighted the high environmental risks due to water column pesticide concentrations in several ponds. Pesticides other than insecticides do not seem to be critical factors to explain the macroinvertebrate assemblages observed here. In contrast to sediment pollution, pollutants measured in the water column provide a snapshot into water quality; sediment pollution is relatively stable over time, and the measurements are more reliable as an indicator of pollution level (Casey et al. 2007; Sun et al. 2019). Conductivity is a general indicator of the presence of many ions in the solution, which is consistent with urban pollution associated with de-icing salts and TE (Brand et al. 2010; Oertli and Parris 2019; Wu et al. 2020). Surprisingly, conductivity is not associated with TE concentrations in our study, suggesting that these two factors do not filter morphotaxa in the same way in different ponds. Conductivity is associated with forest pond J and, to a lesser extent pond G,in addition to Ceratopogoninae, *P. antipodarum* and *P. minutissima*. Though in a rural area, pond J is bordered by a road, which may explain the high conductivity.

Based on morphotaxa abundances, we found two groups of parameters, COD and TE1, in water on the one hand, and fungicide and pharmaceuticals in sediment with dissolved insecticides on the other, generally in the same ponds but at different sampling campaigns. These ponds include G, C, B and E. Ponds B, C and E are located in an agricultural area, ponds C and G are near a farm and pond G is near a medical center. The Chironomidae Orthocladiinae, the mayfly *C. dipterum*, the mollusc *V. macrostoma* and the annelid *Clitellata* sp. characterise the assemblages found in these ponds. These morphotaxa, in particular Chironomidae and annelids, are typical of aquatic systems embedded in a degraded environment (Hill and Wood 2014; Mackintosh et al. 2015; Wood et al. 2001). When considering the PA of morphotaxa, water conductivity is associated with pond J described above.

Compared with the analysis based on abundances, ponds C and G are characterised by high concentrations of pollutants (i.e., pharmaceuticals and fungicides in the sediment). Another set of ponds is associated with high water conductivity and major ion concentrations. This set includes pond J and, to a lesser extent, ponds A, K and L, characterised by the presence of the dipterans Ceratopogoninae, *Chaoborus* sp. and *Chironomus* sp., the Heteroptera *P. minutissima* and the Crustacea *Proasellus* sp. These ponds are mostly surrounded by grasslands and forest, with pond K being the most urbanised pond with nearby dwellings. This points to diffuse pollution associated with road traffic and occasional human activities. In both analyses, the ubiquitous *C. dipterum* is distinguished by its association with high dissolved ion concentrations and, to a lesser extent, by pharmaceuticals in sediment.

Legendre (2014) recommends using PA dissimilarity coefficients when community assemblages are characterised by a high turnover and quantitative indices when the assemblages differ in terms of abundances rather than species diversity. In our study, the high turnover in macroinvertebrate assemblages favours an analysis based on PA, thus concluding the RDA with the least factors: pharmaceuticals sediment and conductivity and ammonium concentration in water. Our third hypothesis is thus partially validated, as contaminants allow us to discriminate several ponds characterised by certain morphotaxa.

### 4.3 Characteristics of the peri-urban environment

The absence of a clear relationship between land use and morphotaxa diversity suggests that the presence of roads, buildings, or impervious surfaces in close vicinity to the ponds is not a critical parameter to explain the observed patterns of morphotaxa diversity at the regional scale. Instead, traces of particular activities influence morphotaxa diversity. For instance, pond A is located in a forest, which explains the hunting cartridges found in and around the pond. We did not find a high level of contaminants in this pond, although its conductivity may be due to the cartridges. Situated between a forest and fields, pond L is not far from residential buildings; people walk to this pond and brush their dogs there (we found a bristle of hairs), also allowing them to swim in the water. In this pond, Nélieu et al. (2020) found high imidacloprid concentrations, which is a veterinary pharmaceutical used to treat dog fleas and ticks. Ponds B and C are both located near farms. We observed that the farmer washed his tractor and equipment in one pond, with the wastewater running off into the pond. Pond G is located near a farm and medical center where carbamazepine, a human anti-epileptic, is used. Pharmaceuticals are markers of human activities, and carbamazepine, which is resistant to biodegradation, is only used by humans (Kasprzyk-Hordern et al. 2009). These observations illustrate the multi-functionality of peri-urban areas with a mixture of different users (Friedmann 2016; Zoomers et al. 2017) at the local scale. Peri-urban ponds combine contaminants typical of rural and urban environments, that is, a runoff of excess nitrogen (and phosphorus) and an influx of heavy metals and salt from road applications (Wanek et al. 2021). To this list, we may add pesticides and pharmaceuticals.

In their review, Oertli and Parris (2019) showed the diversity of criteria used to quantify the urbanisation of a site (presence of buildings, roads, etc.). We suggest here that local and recurrent human actions may blur these categories, particularly in semi-urban areas. Though embedded in a semi-natural matrix, peri-urban ponds are easily reached and may encounter small, chronic and various perturbations. Several authors have advocated for well-managed urban ponds to provide high-quality habitats and support greater biodiversity (Oertli and Parris 2019; Perron et al. 2021). Peri-urban areas tend to be well described, and their potential contribution to sustainable development thus becomes more evident (Wandl and Magoni 2017). As the water quality of peri-urban ponds tends to be more similar to urban ponds than rural ones (Wanek et al. 2021), efforts should be made to manage these systems, especially since they connect rural and urban systems in the blue grid. The use of the LCBD is an interesting approach to identify the ponds to restore (Legendre 2014).

## 5. Conclusion

Our study of macroinvertebrates and water and sediment contaminants in 12 peri-urban ponds over 2 consecutive years reveals a high morphotaxa turnover with the absence of rare and pollutant-sensitive morphotaxa. The macroinvertebrate assemblages were relatively stable, and those contributing the most to regional biodiversity are typical of degraded ponds. The pollutants best describing macroinvertebrate PA in assemblages are pharmaceuticals in sediment and conductivity and ammonium concentration in water. Although an environmental risk due to water column pesticides could be estimated, this factor is not structuring for macroinvertebrate community. Peri-urban areas are characterised by multi-functionality with a mixture of different uses and users. Ponds located in these environments are exposed to various human activities, leading to small, chronic and diverse contaminations that affect macroinvertebrate abundance and community structure.

## Acknowledgments

Sébastien Breuil and Amélie Trouvé (Ecosys Versailles) are thanked for their help in sampling and packaging during all the campaigns and for DOC analysis. Nathalie Bernet (Ecosys Thiverval-Grignon) is acknowledged for PAH analysis. Philippe Beguinel and Henri Roche (CEA) are thanked for their advice in carrying out the project and their participation in the trace analyses. We would like to thank Yves Levi for his support and enriching discussions.

We thank Aurélie Goutte and two anonymous reviewers for their detailed comments, which greatly improved the article. This work was supported by the PSDR4 program.

## Data, scripts, code, and supplementary information availability

All the data and scripts are available online: https://doi.org/10.48579/PRO/TX0PU9. Supplementary information is available online: https://doi.org/10.48579/PRO/TX0PU9. (Not accessible during the peer-review process)

## Supplementary materials

### Appendix 1. Analyses of the sediment

To perform the extraction, duplicate sediment subsamples of 5 g were placed in 50 mL polypropylene tubes (Falcon BD) with a 15 mL mixture of 6/2/2 (v:v) methanol/McIlvaine buffer pH 8/EDTA 0.2 mM being added to each tube. The tubes were shaken on an orbital shaker (10 min, 300 rpm) and sonicated for 30 min before being centrifuged for 10 min at 1,300 g and 4°C. After collecting 10 mL of supernatant, the sediment was again extracted with 10 mL of the same mixture following the same procedure. Then, 10 mL of supernatant was collected and mixed with the first extract. To perform the purification and concentration by solid-phase extraction of the samples, they were then diluted with 100 mL of Milli-Q water, adjusted to pH 7 using 0.5 M NaOH and percolated on HRXA cartridges (500 mg, Macherey-Nagel) preconditioned successively by 5 mL of methanol, acetonitrile and water. After the percolation of the diluted extracts, the cartridges were rinsed with 5 mL of water, dried under vacuum and eluted in a 6 mL mixture of 95:5 (v/v) acetonitrile/formic acid. Samples were then evaporated under a N_2_ stream, dissolved in 3 mL of 8:2 (v/v) water/acetonitrile and analysed before analysis using an ultra-high-performance liquid chromatograph (Acquity UPLC, Waters) coupled through an electrospray interface to a triple quadrupole mass spectrometer (TQD, Waters). Chromatographic separation was carried out on an Acquity BEH C18 column (2.1 mm × 100 mm, 1.7 µm particle size, Waters) at 30°C. The gradient profile started at 95% water with 0.1% acetic acid and linearly progressed to 55% acetonitrile with 0.1% acetic acid at 7.5 min and then 100% at 8.5 min at a flow rate of 0.3 mL/min. After a 4.5 min plateau, the initial conditions were restored and allowed to equilibrate for 2 min. The injection volume was 10 µL. The UHPLC effluent was directed to the electrospray source of the mass spectrometer, operating alternately in positive and negative modes. Capillary voltage (3.0 kV for all the analytes), cone voltage and collision-induced dissociation with argon (under a pressure of 3.4 × 10^−3^ mbar in the collision cell) were optimised individually for each compound to maximise their detection under multiple-reaction monitoring mode (MRM), as indicated in Nélieu et al. (2020). The source temperature was set at 120°C, desolvation temperature at 300°C, extractor voltage at 3 V and cone and desolvation gas flow (nitrogen) at 20 and 800 L/h, respectively (Bourdat-Deschamps et al, 2014).

The external calibration was performed with standard mixtures also analysed by UPLC-MS/MS, with the calibration curves being obtained by weighting 1/x and with residues below 20% using the QuanLynx software (Waters). The matrix effect was corrected by spiking each sample by a known amount of standard mixture in cases where the error exceeded 10%. The quantification limits are given in Table S1.

### Appendix 2. Seasonal or inter-annual effects on morphotaxa abundances

We carried out a correspondence analysis (CA) with the morphotaxa presence/absence. Then we recovered the scores of the ponds on axes 1 and 2 and tested them for season * year effects with an ANOVA (Benedetti et al 2019).

The CA shows that the axes 1 and 2 explain respectively 5.6% and 4.8% of the variance. The ANOVA shows a season effect (p<0.0001) on the CA axis 2. However, as the results are not robust, we conclude that there is no season or year effect on the assemblages.

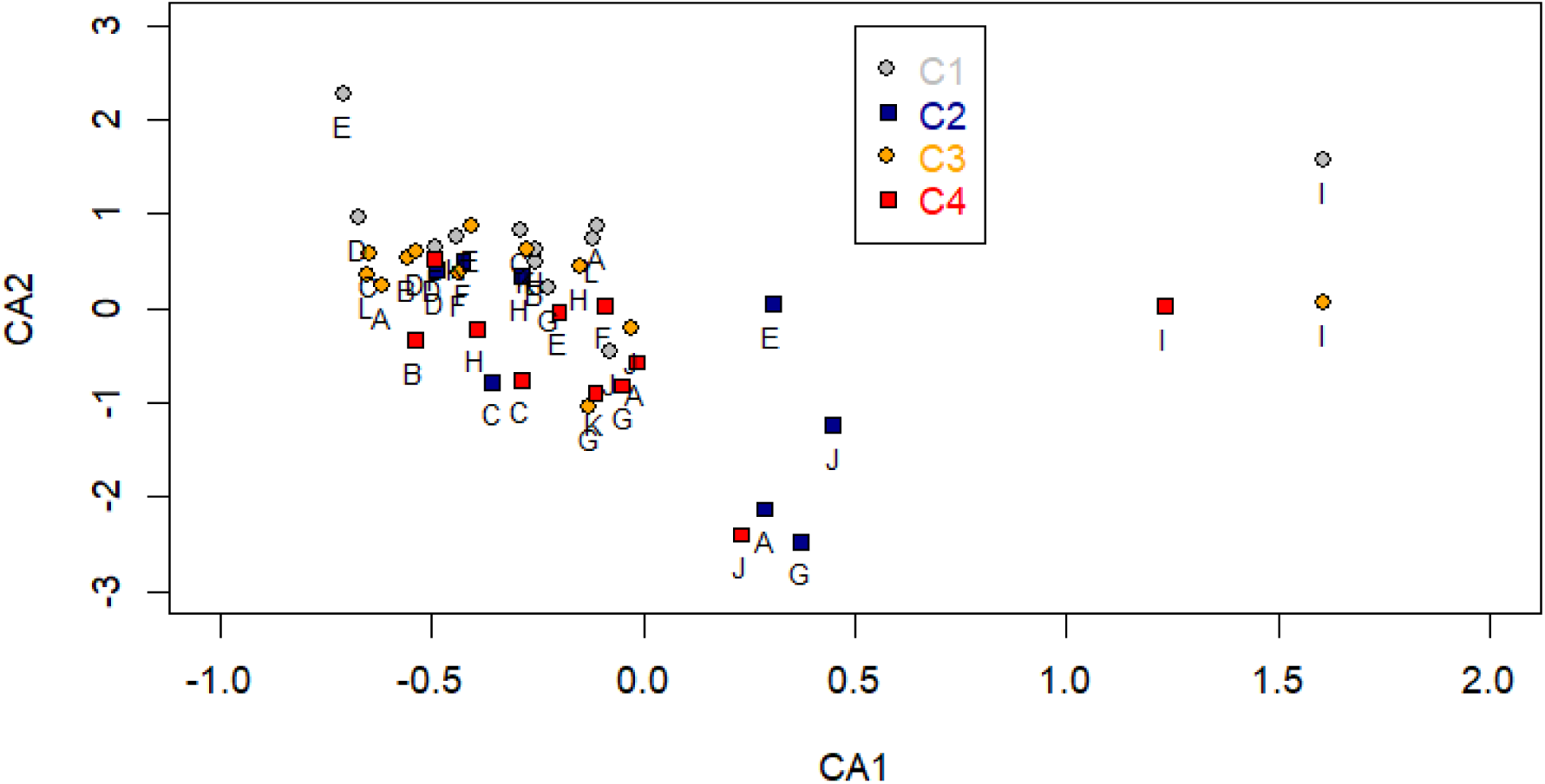

Pond scores on CA axis 1 ∼Season*Year

|  | Df | Sum Sq | Mean Sq | F value | Pr(>F) |
| --- | --- | --- | --- | --- | --- |
| Season | 1 | 1.97 | 1.9733 | 2.128 | 0.152 |
| Year | 1 | 1.16 | 1.1637 | 1.255 | 0.269 |
| Season:Year | 1 | 0.90 | 0.9021 | 0.973 | 0.330 |
| Residuals | 40 | 37.09 | 0.9274 |  |  |

Pond scores on CA axis 2 ∼Season*Year

Pond scores on CA axis 2 ~Season\*Year
|  | Df | Sum Sq | Mean Sq | F value | Pr(>F) |
| --- | --- | --- | --- | --- | --- |
| Season | 1 | 11.434 | 11.434 | 17.637 | **0.000145** |
| Year | 1 | 0.913 | 0.913 | 1.408 | 0.242336 |
| Season:Year | 1 | 0.651 | 0.651 | 1.004 | 0.322414 |
| Residuals | 40 | 25.931 | 0.648 |  |  |

Benedetti et al. 2019. The seasonal and inter-annual fluctuations of plankton abundance and community structure in a North Atlantic marine protected area. Frontiers in Marine Science. doi: 10.3389/fmars.2019.00214

---

**Figure S1.**
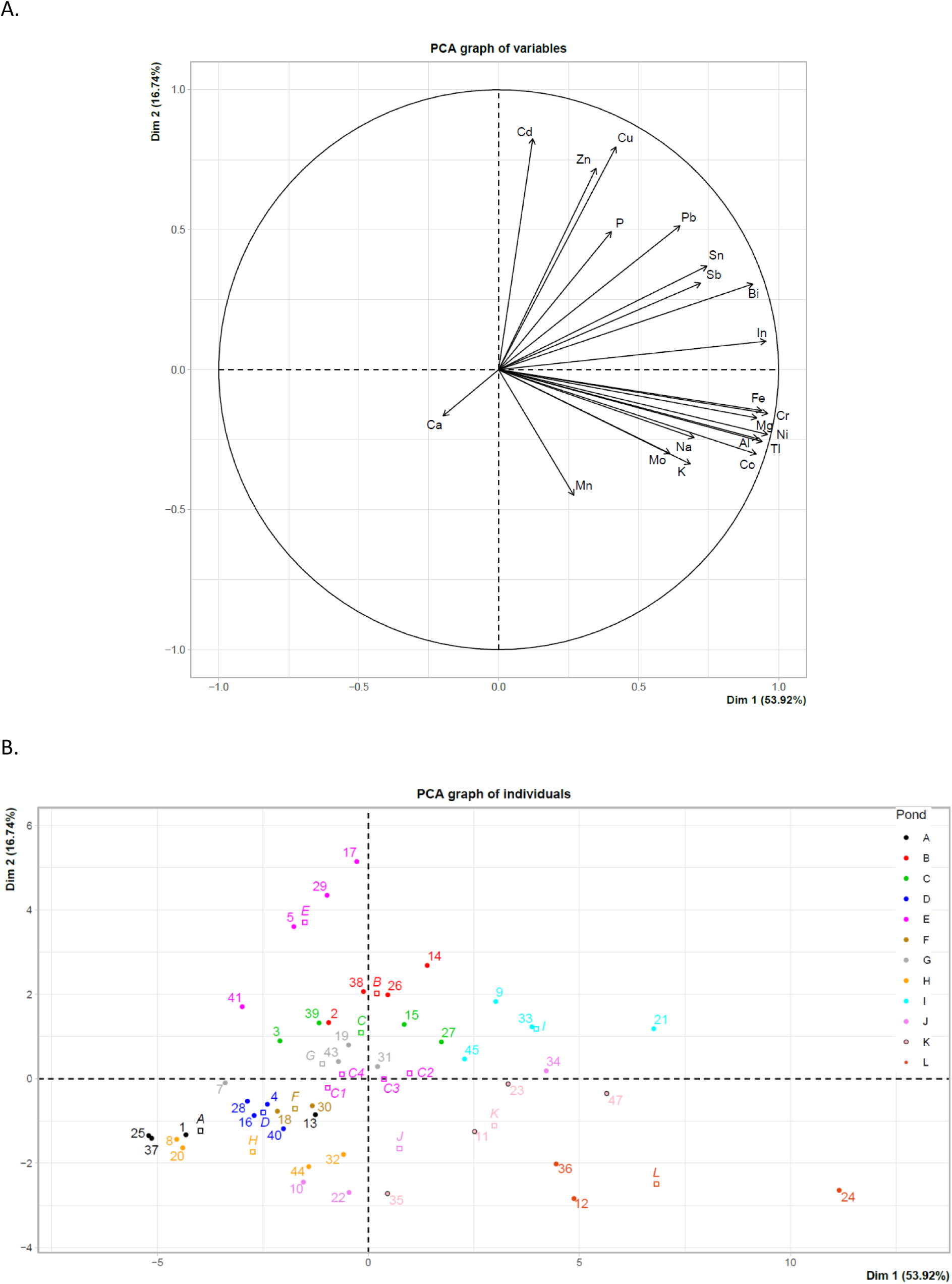
Principal component analysis (PCA) in all ponds and in the four field campaigns with trace elements (TE). A. Correlation circle; B. Graph of the ponds.

**Figure S2.**
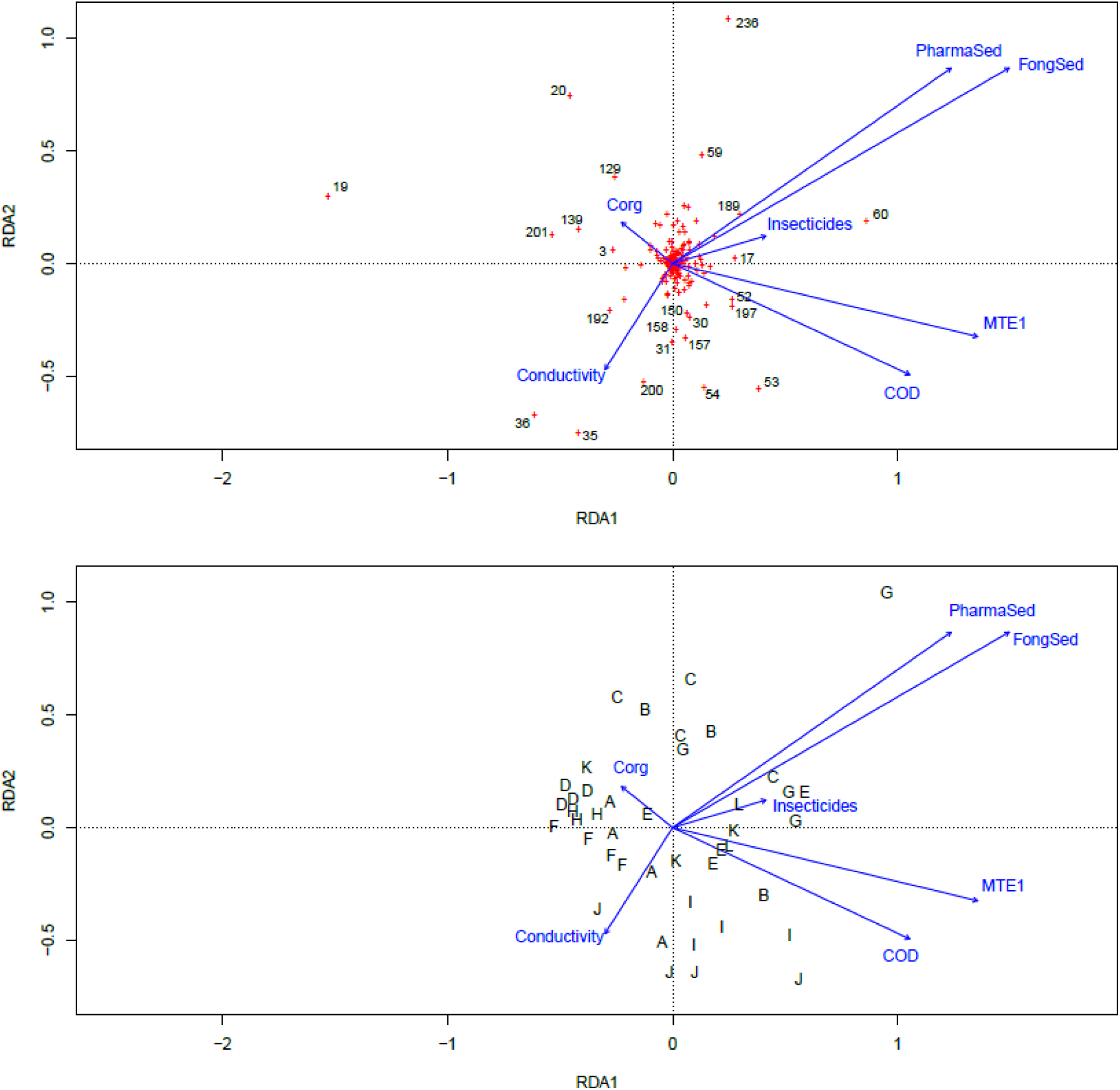
Parsimonious redundancy analysis (RDA) based on morphotaxa abundances. The biplots show the variables (blue arrows) with either the morphotaxa (numbers, panel a) or the ponds (letters, panel b). The number of morphotaxa is shown in brackets: *Acroloxus lacustris* (3); *Asellus* sp. (19); Baetidae 02 (20); Ceratopogoninae 01 (30); Ceratopogoninae 02 (31); *Chaoborus* sp. 01 (35); *Chaoborus* sp. 02 (36); *Chironomus* sp. 01 (53); *Chironomus* sp. 02 (54); Clitellata 01 (59); *Cloeon dipterum* (60); *Helobdella* sp. 02 (129); *Hippeutis complanatus* (139); *Hydrometra stagnorum* (150); *Hygrotus inaequalis* (157); *Hyphydrus ovatus* (158); Orthocladiinae 03 (189), *Physella acuta* (192); *Potamopyrgus antipodarum* (200); *Proasellus* sp. (201); and *Valvata macrostoma* (236). COD: chemical oxygen demand; Conductivity: water conductivity; Corg: organic carbon concentration in water; FongSed: Fungicide concentration in sediment; Insecticides: Insecticide concentration in water; MTE1: Trace element in water – PCA axis 1; PharmaSed: concentration of pharmaceuticals in sediment.

**Table S1.** Exhaustive list of pesticides and pharmaceuticals measured in water and sediments. Only compounds in bold were included in the analysis. The other compounds were not detected, or their values were below the quantification limits.

| Action | Family | Compound |
| --- | --- | --- |
| Herbicide | Triazine and metabolits | **Atrazine** |
|  |  | **Atrazine-desethyl** |
|  |  | **Simazine** |
|  |  | **Terbuthylazine** |
|  |  | **Terbuthylazine-desethyl** |
|  | Isoxazolidinone | **Clomazone** |
|  | Pyridinecarboxamide | **Diflufenican** |
|  | Alkanamide | **Napropamid** |
|  | Chloroacetanilids | **Acetochlore** |
|  |  | **Alachlore** |
|  |  | **Dimethachlore** |
|  |  | **Metolachlore** |
|  | Sulfonylureas | **Chlorsulfuron** |
|  |  | **Metsulfuron-methyl** |
|  |  | **Nicosulfuron** |
|  | Aryloxyalkanoic acid | Mecoprop |
|  | Tricetone | Sulcotrione |
|  |  | Mesotrione |
|  | Hydroxybenzonitriles | Bromoxynil |
|  |  | Ioxynil |
|  | Phenylureas and metabolits | Chlorotoluron |
|  |  | DCPMU |
|  |  | Diuron |
|  |  | Isoproturon |
|  |  | Linuron |
| Safener |  | Dichlormid |
| Fungicide | Triazoles | **Epoiconazole** |
|  |  | **Hexaconazole** |
|  |  | **Metconazole** |
|  |  | **Tebuconazole** |
|  | Carboxamides | **Boscalid** |
|  | Strobilurins | **Picoxystrobine** |
|  |  | **Dimoxystrobine** |
| Insecticide | Neonicotinoids | **Imidaclopride** |
|  | Carbamates | **Pyrimicarbe** |
| Antibiotics | Quinolones | Flumequine |
|  |  | Nalidix Acid |
|  |  | Oxolinic Acid |
|  | Lincosamides | Lincomycine |
|  | Sulfonamides | Sulfadiazine |
|  |  | Sulfamethazine |
|  |  | Sulfamethoxazole |
|  | Tetracycline | Doxycycline |
|  |  | Tetracycline |
| Other pharmaceuticals | Anti-epileptic | **Carbamazepine** |
|  | Anti-inflammatory | Ibuprofen |
|  |  | Diclofenac |

**Table S2.** Effects of the ponds and field sessions on morphotaxa richness, Shannon index and evenness.

| **Model** | **df** | **F-value** | **p-value** |
| --- | --- | --- | --- |
| Morphotaxa richness |  |  |  |
| Pond | 11 | 2.55 | **0.02** |
| Field session | 3 | 6.41 | **0.002** |
| Shannon index |  |  |  |
| Pond | 11 | 1.16 | 0.35 |
| Field session | 3 | 1.11 | 0.36 |
| Evenness |  |  |  |
| Pond | 11 | 1.14 | 0.36 |
| Field session | 3 | 3.79 | **0.02** |

**Table S3.** Total beta diversity and its turnover and nestedness components for the four field sessions computed with the abundance or presence-absence of morphotaxa.

|  | Presence-absence |  |  |  | Abundance |  |  |  |
| --- | --- | --- | --- | --- | --- | --- | --- | --- |
|  | C1 | C3 | C3 | C4 | C1 | C2 | C3 | C4 |
| Total beta diversity | 0.92 | 0.92 | 0.92 | 0.93 | 0.91 | 0.89 | 0.92 | 0.87 |
| Turnover component | 0.90 | 0.86 | 0.86 | 0.90 | 0.85 | 0.80 | 0.86 | 0.78 |
| Nestedness component | 0.02 | 0.06 | 0.06 | 0.03 | 0.06 | 0.09 | 0.06 | 0.8 |

**Table S4.**
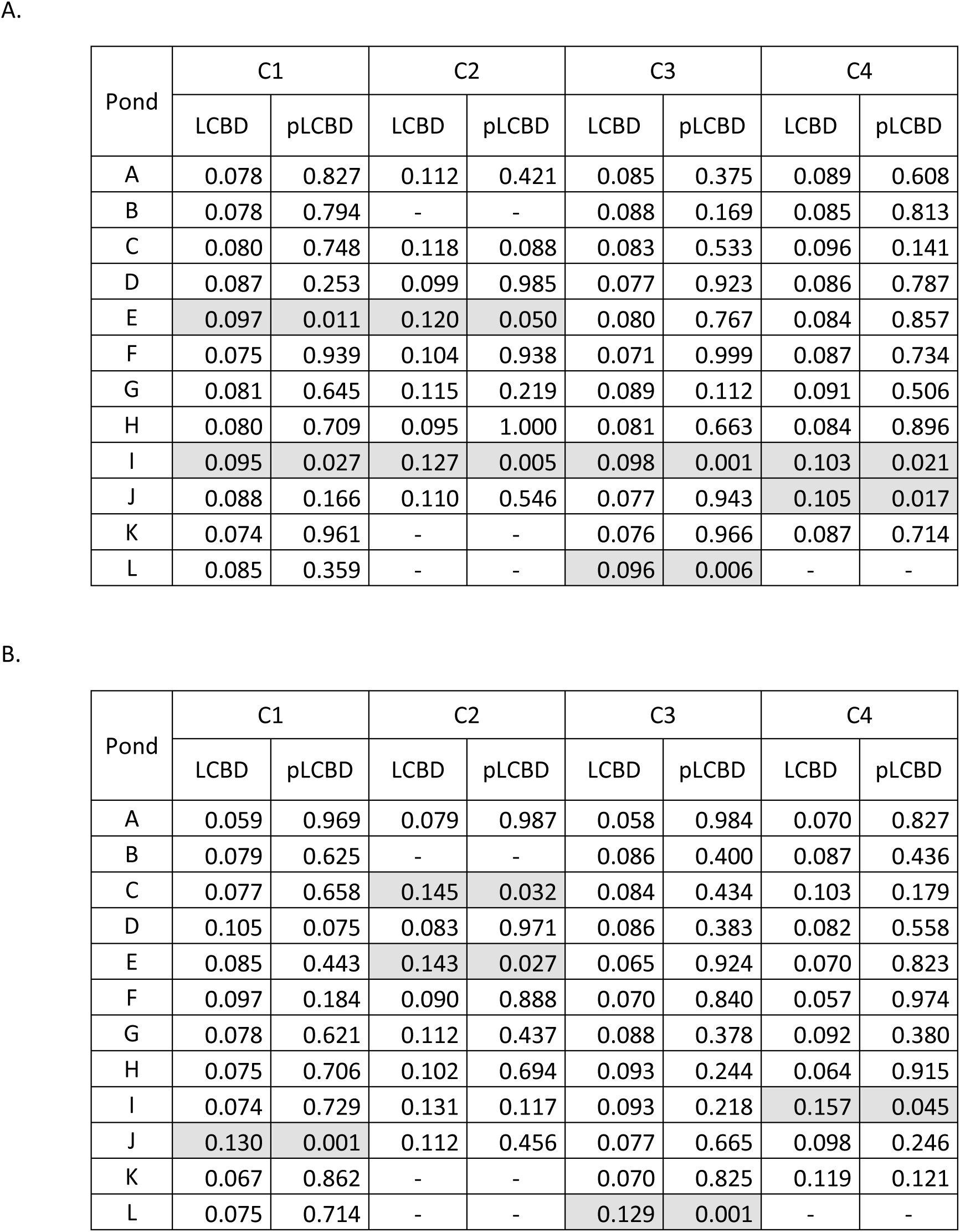
Local contribution to beta diversity (LBCD) of the 12 ponds for the four field sessions and their associated probability for significance. A. LCBD based on morphotaxa presence-absence. B. LCBD based on abundances.

| Pond | C1 |  | C2 |  | C3 |  | C4 |  |
| --- | --- | --- | --- | --- | --- | --- | --- | --- |
|  | LCBD | pLCBD | LCBD | pLCBD | LCBD | pLCBD | LCBD | pLCBD |
| A | 0.078 | 0.827 | 0.112 | 0.421 | 0.085 | 0.375 | 0.089 | 0.608 |
| B | 0.078 | 0.794 | - | - | 0.088 | 0.169 | 0.085 | 0.813 |
| C | 0.080 | 0.748 | 0.118 | 0.088 | 0.083 | 0.533 | 0.096 | 0.141 |
| D | 0.087 | 0.253 | 0.099 | 0.985 | 0.077 | 0.923 | 0.086 | 0.787 |
| E | 0.097 | 0.011 | 0.120 | 0.050 | 0.080 | 0.767 | 0.084 | 0.857 |
| F | 0.075 | 0.939 | 0.104 | 0.938 | 0.071 | 0.999 | 0.087 | 0.734 |
| G | 0.081 | 0.645 | 0.115 | 0.219 | 0.089 | 0.112 | 0.091 | 0.506 |
| H | 0.080 | 0.709 | 0.095 | 1.000 | 0.081 | 0.663 | 0.084 | 0.896 |
| I | 0.095 | 0.027 | 0.127 | 0.005 | 0.098 | 0.001 | 0.103 | 0.021 |
| J | 0.088 | 0.166 | 0.110 | 0.546 | 0.077 | 0.943 | 0.105 | 0.017 |
| K | 0.074 | 0.961 | - | - | 0.076 | 0.966 | 0.087 | 0.714 |
| L | 0.085 | 0.359 | - | - | 0.096 | 0.006 | - | - |

B.
| Pond | C1 |  | C2 |  | C3 |  | C4 |  |
| --- | --- | --- | --- | --- | --- | --- | --- | --- |
|  | LCBD | pLCBD | LCBD | pLCBD | LCBD | pLCBD | LCBD | pLCBD |
| A | 0.059 | 0.969 | 0.079 | 0.987 | 0.058 | 0.984 | 0.070 | 0.827 |
| B | 0.079 | 0.625 | - | - | 0.086 | 0.400 | 0.087 | 0.436 |
| C | 0.077 | 0.658 | 0.145 | 0.032 | 0.084 | 0.434 | 0.103 | 0.179 |
| D | 0.105 | 0.075 | 0.083 | 0.971 | 0.086 | 0.383 | 0.082 | 0.558 |
| E | 0.085 | 0.443 | 0.143 | 0.027 | 0.065 | 0.924 | 0.070 | 0.823 |
| F | 0.097 | 0.184 | 0.090 | 0.888 | 0.070 | 0.840 | 0.057 | 0.974 |
| G | 0.078 | 0.621 | 0.112 | 0.437 | 0.088 | 0.378 | 0.092 | 0.380 |
| H | 0.075 | 0.706 | 0.102 | 0.694 | 0.093 | 0.244 | 0.064 | 0.915 |
| I | 0.074 | 0.729 | 0.131 | 0.117 | 0.093 | 0.218 | 0.157 | 0.045 |
| J | 0.130 | 0.001 | 0.112 | 0.456 | 0.077 | 0.665 | 0.098 | 0.246 |
| K | 0.067 | 0.862 | - | - | 0.070 | 0.825 | 0.119 | 0.121 |
| L | 0.075 | 0.714 | - | - | 0.129 | 0.001 | - | - |

**Table S5.**
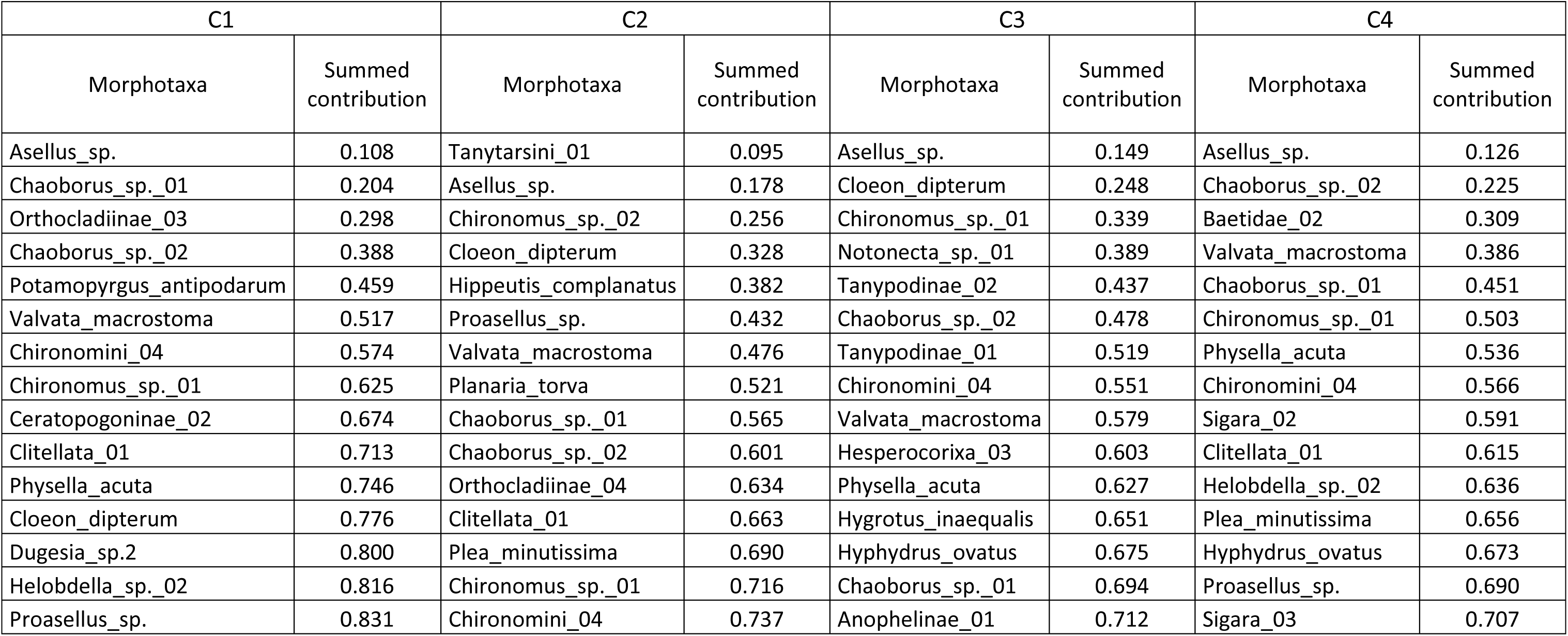
Species contributions to beta diversity (SCBD) for the 15 most common morphotaxa and their summed contribution to the four field sessions. The SCBD calculation is based on morphotaxa abundances.

**Table S6.** Morphotaxa contribution to biodiversity (SCBD) for each field session.

| # | Morphotaxa | C1 | C2 | C3 | C4 |
| --- | --- | --- | --- | --- | --- |
| 1 | *Acilius* sp. | 0.00036 | 0.00000 | 0.00050 | 0.00000 |
| 2 | *Acilius sulcatus* | 0.00000 | 0.00010 | 0.00025 | 0.00042 |
| 3 | *Acroloxus lacustris* | 0.00297 | 0.01012 | 0.00238 | 0.00261 |
| 4 | *Aedes* sp.01 | 0.00000 | 0.00078 | 0.00000 | 0.00090 |
| 5 | *Aeshna cyanea* | 0.00025 | 0.00072 | 0.00230 | 0.00627 |
| 6 | *Aeshna grandis* | 0.00000 | 0.00000 | 0.00010 | 0.00000 |
| 7 | *Aeshna* sp. 01 | 0.00017 | 0.00031 | 0.00297 | 0.00617 |
| 8 | *Aeshna* sp. 02 | 0.00025 | 0.00000 | 0.00000 | 0.00341 |
| 9 | *Aeshna* sp. 03 | 0.00013 | 0.00000 | 0.00036 | 0.00000 |
| 10 | *Aeshna* sp. 04 | 0.00073 | 0.00000 | 0.00609 | 0.00000 |
| 11 | *Aeshna* sp. 05 | 0.00008 | 0.00072 | 0.00014 | 0.00000 |
| 12 | *Aeshna* sp. 06 | 0.00040 | 0.00000 | 0.00260 | 0.00000 |
| 13 | *Aeshna* sp. 07 | 0.00000 | 0.00000 | 0.00516 | 0.00000 |
| 14 | *Alboglossiphonia hyalina* | 0.00000 | 0.00092 | 0.00009 | 0.00158 |
| 15 | *Anacaena* sp. 01 | 0.00000 | 0.00036 | 0.00000 | 0.00000 |
| 16 | *Anacaena* sp. 02 | 0.00057 | 0.00000 | 0.00000 | 0.00000 |
| 17 | *Anophelinae* 01 | 0.00016 | 0.00349 | 0.01782 | 0.00000 |
| 18 | *Armiger crista* | 0.01059 | 0.00889 | 0.00304 | 0.00367 |
| 19 | *Asellus* sp. | 0.10795 | 0.08281 | 0.14862 | 0.12563 |
| 20 | *Baetidae* 02 | 0.01396 | 0.00000 | 0.00000 | 0.08426 |
| 21 | *Batrachobdella* sp. 01 | 0.00061 | 0.00019 | 0.00145 | 0.00440 |
| 22 | *Berosus affinis* | 0.00000 | 0.00076 | 0.00000 | 0.00000 |
| 23 | *Berosus signaticollis* | 0.00000 | 0.00000 | 0.00000 | 0.00042 |
| 24 | *Berosus* sp. 01 | 0.00037 | 0.00000 | 0.00000 | 0.00000 |
| 25 | *Boreobdella* sp. 01 | 0.00009 | 0.00000 | 0.00000 | 0.00000 |
| 26 | *Bythinia* sp. 01 | 0.00017 | 0.00000 | 0.00038 | 0.00141 |
| 27 | *Bythinia* sp. 02 | 0.00000 | 0.00089 | 0.00000 | 0.00000 |
| 28 | *Caenis* sp. 01 | 0.00000 | 0.00000 | 0.00038 | 0.00000 |
| 29 | *Caenis* sp. 02 | 0.00000 | 0.00037 | 0.00000 | 0.00042 |
| 30 | *Ceratopogoninae* 01 | 0.00661 | 0.00703 | 0.00046 | 0.00346 |
| 31 | *Ceratopogoninae* 02 | 0.04913 | 0.00264 | 0.00000 | 0.00042 |
| 32 | *Ceratopogoninae* 03 | 0.00004 | 0.00000 | 0.00000 | 0.00000 |
| 33 | *Ceriagrion* sp 01 | 0.00000 | 0.00000 | 0.00000 | 0.01013 |
| 34 | *Chaoboridae* 01 | 0.00000 | 0.00155 | 0.00000 | 0.00238 |
| 35 | *Chaoborus* sp. 01 | 0.09610 | 0.04427 | 0.01925 | 0.06504 |
| 36 | *Chaoborus* sp. 02 | 0.09000 | 0.03632 | 0.04116 | 0.09934 |
| 37 | *Chaoborus* sp. 03 | 0.00214 | 0.00359 | 0.00185 | 0.00000 |
| 38 | *Chaoborus* sp. 04 | 0.00102 | 0.00082 | 0.00041 | 0.00000 |
| 39 | *Chaoborus* sp. 06 | 0.00015 | 0.00000 | 0.00000 | 0.00000 |
| 40 | *Chironomidae* 01 | 0.00004 | 0.00000 | 0.00000 | 0.00000 |
| 41 | *Chironomidae* 02 | 0.00000 | 0.00100 | 0.00000 | 0.00000 |
| 42 | *Chironomidae* 03 | 0.00000 | 0.00037 | 0.00000 | 0.00000 |
| 43 | *Chironomidae* 04 | 0.00091 | 0.00019 | 0.00000 | 0.00000 |
| 44 | Chironomidae 05 | 0.00009 | 0.00010 | 0.00906 | 0.00000 |
| 45 | Chironomidae 06 | 0.00020 | 0.00000 | 0.00000 | 0.00000 |
| 46 | Chironomidae 07 | 0.00000 | 0.00036 | 0.00000 | 0.00000 |
| 47 | Chironomidae 08 | 0.00000 | 0.00072 | 0.00000 | 0.00000 |
| 48 | Chironomidae 11 | 0.00149 | 0.00000 | 0.00000 | 0.00000 |
| 49 | Chironomini 01 | 0.00112 | 0.00051 | 0.00025 | 0.00000 |
| 50 | Chironomini 02 | 0.00055 | 0.00000 | 0.00000 | 0.00000 |
| 51 | Chironomini 03 | 0.00005 | 0.00000 | 0.00000 | 0.00000 |
| 52 | Chironomini 04 | 0.05723 | 0.02125 | 0.03248 | 0.03042 |
| 53 | *Chironomus* sp. 01 | 0.05097 | 0.02606 | 0.09091 | 0.05198 |
| 54 | *Chironomus* sp. 02 | 0.00000 | 0.07784 | 0.00000 | 0.00000 |
| 55 | *Chironomus* sp. 03 | 0.00059 | 0.00006 | 0.00038 | 0.00000 |
| 56 | *Chironomus* sp. 04 | 0.00000 | 0.00000 | 0.00000 | 0.01219 |
| 57 | *Chironomus* sp. 05 | 0.00105 | 0.00030 | 0.00158 | 0.00000 |
| 58 | *Chironomus* sp. 06 | 0.00009 | 0.00000 | 0.00025 | 0.00000 |
| 59 | Clitellata 01 | 0.03858 | 0.02978 | 0.01273 | 0.02339 |
| 60 | *Cloeon dipterum* | 0.02968 | 0.07220 | 0.09944 | 0.00000 |
| 61 | *Coenagrion scitulum* | 0.00000 | 0.00000 | 0.00000 | 0.00903 |
| 62 | *Coenagrion* sp.01 | 0.00019 | 0.00545 | 0.00507 | 0.01045 |
| 63 | *Coenagrion* sp.02 | 0.00000 | 0.00000 | 0.00000 | 0.00451 |
| 64 | Coenagrionidae 01 | 0.00000 | 0.00074 | 0.00038 | 0.00000 |
| 65 | Coenagrionidae 02 | 0.00000 | 0.00000 | 0.00139 | 0.00000 |
| 66 | Coleoptera 02 | 0.00102 | 0.00000 | 0.00000 | 0.00000 |
| 67 | *Colymbetes fuscus* | 0.00000 | 0.00148 | 0.00124 | 0.00000 |
| 68 | *Colymbetes* sp. 01 | 0.00000 | 0.00000 | 0.00000 | 0.00135 |
| 69 | *Corixa panzeri* | 0.00012 | 0.00000 | 0.00355 | 0.01230 |
| 70 | *Corixa punctata* | 0.00039 | 0.01355 | 0.00426 | 0.00570 |
| 71 | *Corixa* sp.01 | 0.00030 | 0.00201 | 0.00151 | 0.00000 |
| 72 | Corixinae 01 | 0.00063 | 0.00000 | 0.01270 | 0.00000 |
| 73 | Corixinae 02 | 0.01181 | 0.00124 | 0.01261 | 0.00000 |
| 74 | Corixinae 03 | 0.00000 | 0.00021 | 0.00000 | 0.00000 |
| 75 | Crambidae | 0.00000 | 0.00648 | 0.00000 | 0.00000 |
| 76 | *Culex* sp. 01 | 0.00021 | 0.00440 | 0.00078 | 0.00045 |
| 77 | *Culex* sp. 02 | 0.00000 | 0.00648 | 0.00000 | 0.00000 |
| 78 | Culicidae 01 | 0.00000 | 0.00313 | 0.00000 | 0.00000 |
| 79 | Culicidae 02 | 0.00000 | 0.00031 | 0.00000 | 0.00000 |
| 80 | Culicidae 03 | 0.00000 | 0.00062 | 0.00072 | 0.00030 |
| 81 | Culicidae 04 | 0.00000 | 0.00031 | 0.00000 | 0.00000 |
| 82 | Culicidae 05 | 0.00015 | 0.00080 | 0.00142 | 0.00187 |
| 83 | *Cymatia* sp. 01 | 0.00000 | 0.00000 | 0.00072 | 0.00000 |
| 84 | *Cyphon* sp. 01 | 0.00000 | 0.00000 | 0.00036 | 0.00000 |
| 85 | *Cyphon* sp. 02 | 0.00000 | 0.00185 | 0.00000 | 0.00077 |
| 86 | *Cyphon* sp. 03 | 0.00000 | 0.00010 | 0.00000 | 0.00000 |
| 87 | *Dilatata dilatata* | 0.01483 | 0.00817 | 0.00000 | 0.00000 |
| 88 | Diptera 01 | 0.00007 | 0.00000 | 0.00000 | 0.00000 |
| 89 | Diptera 02 | 0.00000 | 0.00036 | 0.00000 | 0.00000 |
| 90 | Diptera 03 | 0.00007 | 0.00000 | 0.00000 | 0.00000 |
| 91 | Diptera 04 | 0.00000 | 0.00000 | 0.00000 | 0.00059 |
| 92 | *Dugesia* sp.1 | 0.00255 | 0.00000 | 0.00114 | 0.00662 |
| 93 | *Dugesia* sp.2 | 0.02393 | 0.00000 | 0.00167 | 0.00000 |
| 94 | Dytiscidae 02 | 0.00000 | 0.00036 | 0.00000 | 0.00000 |
| 95 | *Dytiscus* sp. 01 | 0.00007 | 0.00000 | 0.00047 | 0.00000 |
| 96 | *Enochrus* sp. | 0.00000 | 0.00037 | 0.00000 | 0.00000 |
| 97 | *Erpobdella* sp. 01 | 0.00066 | 0.00000 | 0.00226 | 0.00000 |
| 98 | Erpobdellidae 01 | 0.00063 | 0.00000 | 0.00000 | 0.00000 |
| 99 | *Erythromma viridulum* | 0.00242 | 0.00000 | 0.00326 | 0.01558 |
| 100 | *Ferrissia californica* | 0.00022 | 0.00024 | 0.00000 | 0.00000 |
| 101 | Gammaridae 01 | 0.00305 | 0.00000 | 0.00207 | 0.00000 |
| 102 | Gammaridae 02 | 0.00097 | 0.00000 | 0.00021 | 0.00099 |
| 103 | Gastropoda | 0.00014 | 0.00000 | 0.00000 | 0.00090 |
| 104 | *Gerris lacustris* | 0.00054 | 0.00139 | 0.00046 | 0.00000 |
| 105 | *Gerris* sp.1 | 0.00000 | 0.00000 | 0.00000 | 0.00042 |
| 106 | *Gerris* sp.2 | 0.00004 | 0.00019 | 0.00014 | 0.00000 |
| 107 | *Gerris* sp.3 | 0.00000 | 0.00010 | 0.00087 | 0.00000 |
| 108 | *Gerris* sp.4 | 0.00000 | 0.00000 | 0.00043 | 0.00000 |
| 109 | *Gerris* sp.5 | 0.00000 | 0.00238 | 0.00375 | 0.00000 |
| 110 | *Gerris* sp.6 | 0.00000 | 0.00031 | 0.00000 | 0.00000 |
| 111 | *Gerris* sp.7 | 0.00000 | 0.00145 | 0.00000 | 0.00000 |
| 112 | Glossiphoniidae 01 | 0.00000 | 0.00000 | 0.00000 | 0.00147 |
| 113 | *Gordea* sp. 01 | 0.00024 | 0.00000 | 0.00000 | 0.00000 |
| 114 | *Graphoderus cinereus* | 0.00000 | 0.00021 | 0.00000 | 0.00000 |
| 115 | *Graphoderus* sp. 01 | 0.00006 | 0.00000 | 0.00000 | 0.00000 |
| 116 | *Gyrinus substriatus* | 0.00000 | 0.00168 | 0.00050 | 0.00000 |
| 117 | *Haliphus flavicollis* | 0.00000 | 0.00000 | 0.00000 | 0.00084 |
| 118 | *Haliphus fulvus* | 0.00000 | 0.00031 | 0.00000 | 0.00000 |
| 119 | *Haliphus heydeni* | 0.00028 | 0.00000 | 0.00058 | 0.00699 |
| 120 | *Haliphus immaculatus* | 0.00000 | 0.00010 | 0.00036 | 0.00307 |
| 121 | *Haliphus lineatocollis* | 0.00009 | 0.00455 | 0.00386 | 0.01284 |
| 122 | *Haliphus obliquus* | 0.00000 | 0.00063 | 0.00000 | 0.00506 |
| 123 | *Haliphus ruficollis* | 0.00008 | 0.00031 | 0.00425 | 0.00718 |
| 124 | *Haliphus* sp. 01 | 0.00044 | 0.00000 | 0.00000 | 0.00000 |
| 125 | *Haliphus* sp. 03 | 0.00147 | 0.00058 | 0.00000 | 0.00090 |
| 126 | *Haliphus* sp. 04 | 0.00019 | 0.00060 | 0.00000 | 0.00077 |
| 127 | *Haliphus* sp. 05 | 0.00082 | 0.00036 | 0.00000 | 0.00000 |
| 128 | *Helobdella* sp. 01 | 0.00000 | 0.00037 | 0.00000 | 0.00000 |
| 129 | *Helobdella* sp. 02 | 0.01629 | 0.00457 | 0.00985 | 0.02167 |
| 130 | *Helochares* sp. 01 | 0.00000 | 0.00000 | 0.00068 | 0.00000 |
| 131 | *Helochares* sp. 02 | 0.00019 | 0.00000 | 0.00000 | 0.00000 |
| 132 | *Helochares* sp. 03 | 0.00000 | 0.00037 | 0.00000 | 0.00000 |
| 133 | *Helophorus* sp. 01 | 0.00000 | 0.00000 | 0.00009 | 0.00000 |
| 134 | *Hemiclepsis marginata* | 0.00009 | 0.00000 | 0.00019 | 0.00059 |
| 135 | Hesperocorixa 01 | 0.00011 | 0.00083 | 0.00099 | 0.00587 |
| 136 | Hesperocorixa 02 | 0.00020 | 0.00256 | 0.00109 | 0.00000 |
| 137 | Hesperocorixa 03 | 0.00102 | 0.00368 | 0.02441 | 0.01287 |
| 138 | Hesperocorixa 04 | 0.00000 | 0.00047 | 0.00000 | 0.00859 |
| 139 | *Hippeutis complanatus* | 0.00700 | 0.05408 | 0.00665 | 0.00927 |
| 140 | *Hydra* sp. 01 | 0.00053 | 0.01718 | 0.00000 | 0.00030 |
| 141 | Hydracari 01 | 0.00238 | 0.00000 | 0.00206 | 0.00000 |
| 142 | Hydracari 02 | 0.00000 | 0.00000 | 0.00109 | 0.00000 |
| 143 | Hydracari 03 | 0.00210 | 0.00016 | 0.00178 | 0.00037 |
| 144 | Hydracari 04 | 0.00000 | 0.00208 | 0.00000 | 0.00000 |
| 145 | Hydracari 05 | 0.00000 | 0.00055 | 0.00000 | 0.00000 |
| 146 | Hydracari 06 | 0.00000 | 0.00000 | 0.00000 | 0.00110 |
| 147 | *Hydraena* sp. 01 | 0.00000 | 0.00010 | 0.00000 | 0.00000 |
| 148 | *Hydraena* sp. 02 | 0.00000 | 0.00010 | 0.00000 | 0.00000 |
| 149 | *Hydrobius* sp. 01 | 0.00000 | 0.00072 | 0.00000 | 0.00000 |
| 150 | *Hydrometra stagnorum* | 0.00000 | 0.00036 | 0.01510 | 0.00419 |
| 151 | Hydrophilidae 01 | 0.00004 | 0.00000 | 0.00000 | 0.00000 |
| 152 | Hydroporinae 01 | 0.00013 | 0.00000 | 0.00000 | 0.00000 |
| 153 | *Hydroporus angustatus* | 0.00041 | 0.00216 | 0.00000 | 0.00000 |
| 154 | *Hydroporus figuratus* | 0.00014 | 0.00000 | 0.00000 | 0.00452 |
| 155 | *Hydroporus palustris* | 0.00025 | 0.00024 | 0.00754 | 0.00725 |
| 156 | *Hygrobia hermanni* | 0.00009 | 0.00000 | 0.00000 | 0.00000 |
| 157 | *Hygrotus inaequalis* | 0.00051 | 0.00388 | 0.02378 | 0.01007 |
| 158 | *Hyphydrus ovatus* | 0.00290 | 0.00178 | 0.02369 | 0.01744 |
| 159 | *Ischnura* sp.1 | 0.00284 | 0.00777 | 0.00075 | 0.00591 |
| 160 | *Ischnura* sp.2 | 0.00000 | 0.00163 | 0.00038 | 0.00000 |
| 161 | *Ischnura* sp.3 | 0.00000 | 0.00037 | 0.00056 | 0.00000 |
| 162 | *Laccobius* sp. 01 | 0.00000 | 0.00000 | 0.00038 | 0.00000 |
| 163 | *Laccophilus minutus* | 0.00000 | 0.00530 | 0.00000 | 0.00147 |
| 164 | *Laccophilus* sp. 01 | 0.00130 | 0.00000 | 0.00053 | 0.00000 |
| 165 | *Laccophilus* sp. 02 | 0.00206 | 0.00000 | 0.00000 | 0.00000 |
| 166 | *Laccophilus* sp. 03 | 0.00188 | 0.00000 | 0.00000 | 0.00000 |
| 167 | *Lestes viridis* | 0.00105 | 0.00050 | 0.00924 | 0.00000 |
| 168 | Libellulidae 01 | 0.00000 | 0.00000 | 0.00000 | 0.00037 |
| 169 | Limnephilidae 02 | 0.00000 | 0.00000 | 0.00000 | 0.00045 |
| 170 | *Limnoxenus niger* | 0.00018 | 0.00000 | 0.00000 | 0.00000 |
| 171 | *Micronecta* sp. 01 | 0.00017 | 0.00000 | 0.00036 | 0.00000 |
| 172 | *Microvelia* sp. 01 | 0.00009 | 0.00245 | 0.00000 | 0.00042 |
| 173 | *Microvelia* sp. 02 | 0.00000 | 0.00000 | 0.00046 | 0.00000 |
| 174 | *Musculium* sp.2 | 0.00044 | 0.00000 | 0.00000 | 0.00000 |
| 175 | *Musculium* sp.3 | 0.00056 | 0.00000 | 0.00920 | 0.00118 |
| 176 | *Myxas* sp. | 0.00287 | 0.00000 | 0.00000 | 0.00000 |
| 177 | *Naucoris maculatus* | 0.00057 | 0.00000 | 0.00449 | 0.00000 |
| 178 | *Noterus clavicornis* | 0.00000 | 0.00000 | 0.00036 | 0.00042 |
| 179 | *Notonecta* sp. 01 | 0.00030 | 0.00519 | 0.05028 | 0.00833 |
| 180 | *Notonecta* sp. 02 | 0.00666 | 0.00000 | 0.00869 | 0.00000 |
| 181 | *Ochthebius minutus* | 0.00000 | 0.00016 | 0.00000 | 0.00045 |
| 182 | *Odontomyia* sp.01 | 0.00008 | 0.00000 | 0.00000 | 0.00000 |
| 183 | *Odontomyia* sp.02 | 0.00004 | 0.00000 | 0.00000 | 0.00000 |
| 184 | *Oplodontha* sp. 01 | 0.00000 | 0.00037 | 0.00000 | 0.00000 |
| 185 | *Oplodontha* sp. 02 | 0.00057 | 0.00000 | 0.00000 | 0.00000 |
| 186 | *Orthetrum* sp. 01 | 0.00000 | 0.00016 | 0.00000 | 0.00000 |
| 187 | Orthoclaadiinae 01 | 0.00575 | 0.00000 | 0.00011 | 0.00000 |
| 188 | Orthoclaadiinae 02 | 0.00632 | 0.00290 | 0.00000 | 0.00000 |
| 189 | Orthoclaadiinae 03 | 0.09402 | 0.00178 | 0.00767 | 0.00000 |
| 190 | Orthoclaadiinae 04 | 0.00209 | 0.03250 | 0.00000 | 0.00000 |
| 191 | *Peltodytes caesus* | 0.00004 | 0.00000 | 0.00000 | 0.00042 |
| 192 | *Physella acuta* | 0.03366 | 0.00443 | 0.02411 | 0.03215 |
| 193 | *Pisidium* sp.1 | 0.00068 | 0.00496 | 0.01413 | 0.00000 |
| 194 | *Pisidium* sp.2 | 0.00000 | 0.00000 | 0.00000 | 0.00074 |
| 195 | *Planaria torva* | 0.00020 | 0.04437 | 0.00000 | 0.00000 |
| 196 | *Platycnemis* sp. 01 | 0.00000 | 0.00412 | 0.00000 | 0.00223 |
| 197 | *Plea minutissima* | 0.00131 | 0.02655 | 0.01421 | 0.01942 |
| 198 | *Podura aquatica* | 0.00000 | 0.00000 | 0.00210 | 0.00202 |
| 199 | Polycentropodidae 01 | 0.00007 | 0.00000 | 0.00000 | 0.00000 |
| 200 | *Potamopyrgus antipodarum* | 0.07138 | 0.00256 | 0.00589 | 0.00539 |
| 201 | *Proasellus* sp. | 0.01505 | 0.04956 | 0.00906 | 0.01720 |
| 202 | *Procambarus clarkii* | 0.00000 | 0.00000 | 0.00010 | 0.00000 |
| 203 | *Pyrrhosoma nymphula* | 0.00000 | 0.00268 | 0.00046 | 0.00000 |
| 204 | *Ranatra linearis* | 0.00000 | 0.00000 | 0.00038 | 0.00000 |
| 205 | *Rhantus suturalis* | 0.00000 | 0.00000 | 0.00000 | 0.00090 |
| 206 | Sciomyzidae 01 | 0.00006 | 0.00000 | 0.00000 | 0.00000 |
| 207 | *Segmentina nitida* | 0.00199 | 0.00000 | 0.00000 | 0.00000 |
| 208 | *Sialis* sp. 01 | 0.00017 | 0.00013 | 0.00077 | 0.00000 |
| 209 | Sigara 01 | 0.00027 | 0.00041 | 0.00000 | 0.00119 |
| 210 | Sigara 02 | 0.00000 | 0.00310 | 0.00000 | 0.02538 |
| 211 | Sigara 03 | 0.00182 | 0.00219 | 0.00000 | 0.01701 |
| 212 | Sigara 04 | 0.00028 | 0.00047 | 0.00000 | 0.00000 |
| 213 | Sigara 06 | 0.00000 | 0.00221 | 0.00318 | 0.00000 |
| 214 | *Sympetrum* sp.1 | 0.00000 | 0.00000 | 0.00050 | 0.00000 |
| 215 | *Sympetrum* sp.2 | 0.00000 | 0.00000 | 0.00093 | 0.00000 |
| 216 | *Sympetrum* sp.3 | 0.00297 | 0.00000 | 0.01068 | 0.00000 |
| 217 | Syrphidae 01 | 0.00000 | 0.00000 | 0.00028 | 0.00000 |
| 218 | Tabanidae 01 | 0.00000 | 0.00010 | 0.00000 | 0.00000 |
| 219 | Tabanidae 02 | 0.00000 | 0.00021 | 0.00000 | 0.00000 |
| 220 | Tanypodinae 01 | 0.00040 | 0.00000 | 0.04110 | 0.00000 |
| 221 | Tanypodinae 02 | 0.00383 | 0.00110 | 0.04738 | 0.00000 |
| 222 | Tanypodinae 03 | 0.00150 | 0.00125 | 0.00000 | 0.00000 |
| 223 | Tanypodinae 04 | 0.00074 | 0.00978 | 0.00207 | 0.00255 |
| 224 | Tanypodinae 05 | 0.00000 | 0.01153 | 0.00000 | 0.00000 |
| 225 | Tanypodinae 06 | 0.00000 | 0.00733 | 0.00000 | 0.01252 |
| 226 | Tanypodinae 07 | 0.00020 | 0.00072 | 0.00000 | 0.00000 |
| 227 | Tanypodinae 08 | 0.00000 | 0.00031 | 0.00000 | 0.00000 |
| 228 | Tanypodinae 09 | 0.00000 | 0.00254 | 0.00452 | 0.00000 |
| 229 | Tanypodinae 10 | 0.00000 | 0.00691 | 0.00000 | 0.00000 |
| 230 | Tanypodinae 11 | 0.00000 | 0.00000 | 0.00000 | 0.00845 |
| 231 | *Tanysphyrus lemnae* | 0.00000 | 0.00108 | 0.00000 | 0.00000 |
| 232 | Tanytarsini 01 | 0.00320 | 0.09510 | 0.00000 | 0.00000 |
| 233 | Tanytarsini 02 | 0.00182 | 0.00000 | 0.00000 | 0.00000 |
| 234 | *Theremyzon* sp. 01 | 0.00034 | 0.00016 | 0.00029 | 0.00000 |
| 235 | Trichoptera 02 | 0.00000 | 0.00000 | 0.00000 | 0.00089 |
| 236 | *Valvata macrostoma* | 0.05748 | 0.04464 | 0.02756 | 0.07712 |

**Table S7.** Results of the redundancy analyses (RDA) parsimonious models based on morphotaxa abundance.

| **Variables** | **Cumulative adjusted R<sup>2</sup>** | **F value** | ***p* value** |
| --- | --- | --- | --- |
| Fungicide concentration in sediment | 0.047 | 3.142 | 0.001 |
| MTE1 | 0.079 | 2.465 | 0.002 |
| Water conductivity | 0.098 | 1.862 | 0.014 |
| COD | 0.115 | 1.737 | 0.024 |
| Pharmaceutical concentration in sediment | 0.131 | 1.748 | 0.022 |
| Insecticide concentration in water | 0.146 | 1.644 | 0.034 |
| Organic C concentration in water | 0.160 | 1.601 | 0.05 |

**Table S8.** Morphotaxa scores in the presence-absence-based redundancy analysis (RDA) and the abundance-based RDA.

| # | Morphotaxa | Presence-absence-based RDA |  | Abundance-based RDA |  |
| --- | --- | --- | --- | --- | --- |
|  |  | RDA1 | RDA2 | RDA1 | RDA2 |
| 1 | *Acilius* sp. | -0.006 | 0.002 | 0.001 | 0.001 |
| 2 | *Acilius sulcatus* | -0.007 | -0.016 | -0.005 | -0.013 |
| 3 | *Acroloxus lacustris* | -0.061 | -0.055 | -0.112 | 0.024 |
| 4 | *Aedes* sp.01 | -0.009 | -0.011 | -0.003 | -0.012 |
| 5 | *Aeshna cyanea* | 0.043 | 0.077 | 0.029 | -0.026 |
| 6 | *Aeshna grandis* | -0.002 | -0.003 | -0.002 | -0.001 |
| 7 | *Aeshna* sp. 01 | -0.004 | -0.003 | 0.012 | -0.056 |
| 8 | *Aeshna* sp. 02 | 0.019 | 0.027 | 0.002 | 0.012 |
| 9 | *Aeshna* sp. 03 | 0.023 | -0.009 | 0.009 | 0.019 |
| 10 | *Aeshna* sp. 04 | -0.009 | 0.033 | -0.029 | 0.022 |
| 11 | *Aeshna* sp. 05 | -0.005 | 0.003 | 0.006 | -0.015 |
| 12 | *Aeshna* sp. 06 | 0.032 | 0.026 | 0.023 | 0.005 |
| 13 | *Aeshna* sp. 07 | -0.010 | 0.021 | 0.022 | -0.052 |
| 14 | *Alboglossiphonia hyalina* | -0.016 | -0.015 | -0.028 | 0.009 |
| 15 | *Anacaena* sp. 01 | 0.000 | 0.008 | 0.005 | -0.008 |
| 16 | *Anacaena* sp. 02 | -0.004 | -0.003 | 0.003 | -0.003 |
| 17 | *Anophelinae* 01 | 0.095 | -0.152 | 0.117 | 0.008 |
| 18 | *Armiger crista* | -0.063 | 0.013 | -0.088 | -0.009 |
| 19 | *Asellus* sp. | -0.416 | -0.198 | -0.648 | 0.124 |
| 20 | *Baetidae* 02 | -0.066 | 0.194 | -0.193 | 0.313 |
| 21 | *Batrachobdella* sp. 01 | 0.070 | -0.016 | 0.020 | 0.069 |
| 22 | *Berosus affinis* | 0.009 | -0.004 | 0.006 | 0.010 |
| 23 | *Berosus signaticollis* | -0.005 | -0.014 | -0.004 | -0.006 |
| 24 | *Berosus* sp. 01 | -0.001 | 0.008 | 0.003 | -0.001 |
| 25 | *Boreobdella* sp. 01 | -0.002 | -0.003 | 0.001 | 0.004 |
| 26 | *Bythinia* sp. 01 | -0.009 | -0.005 | -0.017 | 0.010 |
| 27 | *Bythinia* sp. 02 | -0.004 | -0.004 | -0.008 | 0.003 |
| 28 | *Caenis* sp. 01 | -0.002 | 0.002 | 0.003 | -0.002 |
| 29 | *Caenis* sp. 02 | 0.002 | -0.024 | 0.002 | -0.004 |
| 30 | *Ceratopogoninae* 01 | -0.044 | 0.000 | 0.032 | -0.102 |
| 31 | *Ceratopogoninae* 02 | -0.087 | -0.257 | -0.001 | -0.150 |
| 32 | *Ceratopogoninae* 03 | -0.002 | -0.005 | 0.000 | -0.004 |
| 33 | *Ceriagrion* sp 01 | -0.023 | -0.070 | -0.019 | -0.030 |
| 34 | *Chaoboridae* 01 | -0.014 | -0.035 | -0.014 | -0.023 |
| 35 | *Chaoborus* sp. 01 | -0.168 | 0.063 | -0.177 | -0.319 |
| 36 | *Chaoborus* sp. 02 | -0.220 | 0.014 | -0.259 | -0.287 |
| 37 | *Chaoborus* sp. 03 | -0.036 | -0.056 | -0.009 | -0.061 |
| 38 | *Chaoborus* sp. 04 | -0.006 | 0.020 | -0.010 | 0.001 |
| 39 | *Chaoborus* sp. 06 | -0.003 | -0.010 | 0.000 | -0.008 |
| 40 | *Chironomidae* 01 | 0.004 | 0.001 | 0.000 | 0.004 |
| 41 | Chironomidae 02 | 0.003 | 0.007 | 0.001 | 0.010 |
| 42 | Chironomidae 03 | 0.007 | -0.009 | 0.005 | 0.003 |
| 43 | Chironomidae 04 | -0.006 | -0.008 | -0.013 | 0.001 |
| 44 | Chironomidae 05 | -0.010 | 0.054 | 0.030 | -0.006 |
| 45 | Chironomidae 06 | 0.000 | 0.009 | 0.001 | -0.004 |
| 46 | Chironomidae 07 | 0.000 | 0.008 | 0.005 | -0.008 |
| 47 | Chironomidae 08 | 0.000 | 0.012 | 0.008 | -0.011 |
| 48 | Chironomidae 11 | 0.019 | -0.036 | 0.011 | 0.002 |
| 49 | Chironomini 01 | -0.009 | -0.026 | -0.005 | 0.005 |
| 50 | Chironomini 02 | -0.001 | 0.010 | 0.004 | -0.002 |
| 51 | Chironomini 03 | -0.001 | -0.001 | -0.002 | 0.000 |
| 52 | Chironomini 04 | -0.011 | 0.199 | 0.112 | -0.070 |
| 53 | *Chironomus* sp. 01 | -0.076 | 0.161 | 0.162 | -0.236 |
| 54 | *Chironomus* sp. 02 | -0.069 | -0.156 | 0.059 | -0.234 |
| 55 | *Chironomus* sp. 03 | -0.003 | -0.002 | 0.004 | -0.001 |
| 56 | *Chironomus* sp. 04 | -0.013 | 0.033 | 0.000 | 0.002 |
| 57 | *Chironomus* sp. 05 | -0.014 | 0.009 | -0.002 | 0.010 |
| 58 | *Chironomus* sp. 06 | -0.004 | -0.001 | 0.000 | 0.001 |
| 59 | Clitellata 01 | 0.147 | 0.089 | 0.055 | 0.204 |
| 60 | *Cloeon dipterum* | 0.453 | -0.401 | 0.365 | 0.078 |
| 61 | *Coenagrion scitulum* | 0.040 | 0.028 | -0.002 | 0.039 |
| 62 | *Coenagrion* sp.01 | 0.122 | -0.080 | 0.045 | 0.078 |
| 63 | *Coenagrion* sp.02 | 0.030 | 0.018 | -0.001 | 0.029 |
| 64 | Coenagrionidae 01 | 0.007 | -0.011 | 0.011 | 0.002 |
| 65 | Coenagrionidae 02 | -0.008 | -0.019 | 0.013 | -0.024 |
| 66 | Coleoptera 02 | -0.001 | 0.021 | 0.002 | -0.010 |
| 67 | *Colymbetes fuscus* | 0.009 | -0.015 | 0.009 | -0.002 |
| 68 | *Colymbetes* sp. 01 | -0.002 | 0.014 | -0.006 | 0.004 |
| 69 | *Corixa panzeri* | 0.084 | 0.043 | 0.022 | 0.106 |
| 70 | *Corixa punctata* | 0.015 | 0.080 | 0.010 | 0.079 |
| 71 | *Corixa* sp.01 | 0.026 | 0.021 | 0.001 | 0.022 |
| 72 | Corixinae 01 | -0.015 | 0.042 | -0.041 | 0.023 |
| 73 | Corixinae 02 | -0.004 | 0.019 | 0.050 | 0.013 |
| 74 | Corixinae 03 | -0.001 | -0.004 | 0.000 | -0.006 |
| 75 | Crambidae | -0.001 | 0.035 | 0.023 | -0.032 |
| 76 | *Culex* sp. 01 | 0.012 | -0.087 | 0.007 | -0.012 |
| 77 | *Culex* sp. 02 | -0.021 | -0.069 | 0.004 | -0.047 |
| 78 | Culicidae 01 | -0.014 | -0.046 | 0.003 | -0.030 |
| 79 | Culicidae 02 | -0.004 | -0.015 | 0.001 | -0.009 |
| 80 | Culicidae 03 | 0.031 | -0.020 | 0.013 | 0.017 |
| 81 | Culicidae 04 | -0.001 | -0.005 | 0.000 | -0.007 |
| 82 | Culicidae 05 | 0.029 | -0.060 | 0.021 | -0.001 |
| 83 | *Cymatia* sp. 01 | 0.034 | -0.012 | 0.016 | 0.027 |
| 84 | *Cyphon* sp. 01 | 0.024 | -0.008 | 0.012 | 0.019 |
| 85 | *Cyphon* sp. 02 | 0.013 | -0.009 | 0.016 | 0.005 |
| 86 | *Cyphon* sp. 03 | 0.000 | -0.003 | 0.000 | -0.004 |
| 87 | *Dilatata dilatata* | -0.031 | -0.040 | -0.059 | -0.003 |
| 88 | Diptera 01 | -0.001 | -0.001 | -0.002 | 0.000 |
| 89 | Diptera 02 | 0.000 | 0.008 | 0.005 | -0.008 |
| 90 | Diptera 03 | -0.001 | -0.001 | -0.002 | 0.000 |
| 91 | Diptera 04 | -0.003 | -0.002 | -0.005 | 0.001 |
| 92 | *Dugesia* sp.1 | 0.067 | 0.017 | -0.010 | 0.091 |
| 93 | *Dugesia* sp.2 | 0.145 | -0.005 | 0.029 | 0.105 |
| 94 | Dytiscidae 02 | 0.000 | 0.008 | 0.005 | -0.008 |
| 95 | *Dytiscus* sp. 01 | -0.002 | 0.009 | -0.007 | 0.005 |
| 96 | *Enochrus* sp. | 0.007 | -0.009 | 0.005 | 0.003 |
| 97 | *Erpobdella* sp. 01 | -0.011 | 0.007 | 0.015 | -0.001 |
| 98 | Erpobdellidae 01 | -0.005 | -0.006 | 0.004 | 0.001 |
| 99 | *Erythromma viridulum* | 0.094 | -0.144 | 0.031 | 0.041 |
| 100 | *Ferrissia californica* | -0.006 | -0.011 | -0.008 | -0.004 |
| 101 | Gammaridae 01 | -0.013 | -0.002 | -0.030 | 0.015 |
| 102 | Gammaridae 02 | -0.014 | -0.010 | -0.023 | 0.003 |
| 103 | Gastropoda | -0.003 | 0.010 | -0.007 | 0.003 |
| 104 | *Gerris lacustris* | -0.016 | -0.027 | -0.004 | -0.025 |
| 105 | *Gerris* sp.1 | -0.005 | -0.014 | -0.004 | -0.006 |
| 106 | *Gerris* sp.2 | -0.005 | -0.008 | -0.006 | -0.002 |
| 107 | *Gerris* sp.3 | -0.008 | -0.007 | 0.000 | -0.014 |
| 108 | *Gerris* sp.4 | 0.022 | -0.011 | 0.010 | 0.018 |
| 109 | *Gerris* sp.5 | -0.009 | -0.008 | 0.008 | -0.027 |
| 110 | *Gerris* sp.6 | -0.001 | -0.005 | 0.000 | -0.007 |
| 111 | *Gerris* sp.7 | -0.001 | -0.011 | -0.001 | -0.016 |
| 112 | Glossiphoniidae 01 | 0.017 | 0.006 | 0.001 | 0.012 |
| 113 | *Gordea* sp. 01 | -0.002 | 0.005 | 0.001 | -0.001 |
| 114 | *Graphoderus cinereus* | -0.001 | -0.004 | 0.000 | -0.006 |
| 115 | *Graphoderus* sp. 01 | 0.000 | 0.003 | 0.000 | -0.001 |
| 116 | *Gyrinus substriatus* | -0.008 | -0.004 | -0.011 | -0.002 |
| 117 | *Haliphus flavicollis* | -0.007 | -0.020 | -0.005 | -0.009 |
| 118 | *Haliphus fulvus* | -0.004 | -0.015 | 0.001 | -0.009 |
| 119 | *Haliphus heydeni* | -0.005 | 0.031 | 0.004 | -0.006 |
| 120 | *Haliphus immaculatus* | 0.021 | 0.012 | 0.020 | 0.014 |
| 121 | *Haliphus lineatocollis* | 0.059 | -0.050 | 0.021 | 0.036 |
| 122 | *Haliphus obliquus* | -0.023 | -0.070 | -0.012 | -0.035 |
| 123 | *Haliphus ruficollis* | -0.039 | -0.084 | -0.009 | -0.059 |
| 124 | *Haliphus* sp. 01 | 0.008 | 0.010 | 0.000 | 0.005 |
| 125 | *Haliphus* sp. 03 | -0.016 | -0.010 | 0.002 | 0.002 |
| 126 | *Haliphus* sp. 04 | -0.006 | -0.008 | 0.005 | -0.012 |
| 127 | *Haliphus* sp. 05 | 0.021 | -0.015 | 0.015 | 0.001 |
| 128 | *Helobdella* sp. 01 | -0.003 | -0.003 | -0.005 | 0.001 |
| 129 | *Helobdella* sp. 02 | 0.007 | 0.097 | -0.109 | 0.160 |
| 130 | *Helochares* sp. 01 | -0.001 | 0.012 | 0.005 | 0.003 |
| 131 | *Helochares* sp. 02 | 0.007 | -0.012 | 0.005 | 0.001 |
| 132 | *Helochares* sp. 03 | 0.007 | -0.009 | 0.005 | 0.003 |
| 133 | *Helophorus* sp. 01 | -0.001 | -0.001 | -0.003 | 0.001 |
| 134 | *Hemiclepsis marginata* | 0.000 | -0.012 | -0.005 | 0.003 |
| 135 | *Hesperocorixa* 01 | 0.007 | -0.003 | -0.008 | 0.007 |
| 136 | *Hesperocorixa* 02 | 0.057 | -0.043 | 0.031 | 0.036 |
| 137 | *Hesperocorixa* 03 | -0.118 | -0.097 | -0.091 | -0.068 |
| 138 | *Hesperocorixa* 04 | -0.019 | -0.034 | -0.012 | -0.022 |
| 139 | *Hippeutis complanatus* | -0.088 | -0.064 | -0.177 | 0.063 |
| 140 | *Hydra* sp. 01 | -0.017 | -0.006 | -0.042 | 0.033 |
| 141 | *Hydracari* 01 | 0.002 | 0.040 | -0.013 | 0.025 |
| 142 | *Hydracari* 02 | -0.009 | -0.014 | 0.010 | -0.023 |
| 143 | *Hydracari* 03 | 0.084 | 0.000 | 0.023 | 0.058 |
| 144 | *Hydracari* 04 | 0.002 | 0.007 | -0.002 | 0.014 |
| 145 | *Hydracari* 05 | 0.002 | 0.002 | 0.000 | 0.003 |
| 146 | *Hydracari* 06 | -0.002 | 0.020 | 0.001 | 0.002 |
| 147 | *Hydraena* sp. 01 | 0.000 | -0.003 | 0.000 | -0.004 |
| 148 | *Hydraena* sp. 02 | 0.000 | -0.003 | 0.000 | -0.004 |
| 149 | *Hydrobius* sp. 01 | 0.000 | 0.012 | 0.008 | -0.011 |
| 150 | *Hydrometra stagnorum* | -0.014 | 0.102 | 0.027 | -0.096 |
| 151 | Hydrophilidae 01 | -0.002 | -0.005 | 0.000 | -0.004 |
| 152 | Hydroporinae 01 | -0.003 | -0.004 | -0.001 | 0.004 |
| 153 | *Hydroporus angustatus* | -0.001 | 0.034 | 0.014 | -0.025 |
| 154 | *Hydroporus figuratus* | -0.005 | 0.025 | -0.012 | 0.006 |
| 155 | *Hydroporus palustris* | -0.035 | 0.048 | -0.020 | -0.036 |
| 156 | *Hygrobia hermanni* | 0.005 | -0.009 | 0.003 | 0.000 |
| 157 | *Hygrotus inaequalis* | -0.018 | 0.158 | 0.024 | -0.140 |
| 158 | *Hyphydrus ovatus* | -0.063 | 0.098 | 0.006 | -0.125 |
| 159 | *Ischnura* sp.1 | 0.041 | -0.045 | 0.032 | 0.025 |
| 160 | *Ischnura* sp.2 | -0.010 | 0.000 | 0.010 | 0.005 |
| 161 | *Ischnura* sp.3 | 0.017 | 0.001 | 0.003 | 0.004 |
| 162 | *Laccobius* sp. 01 | -0.002 | 0.002 | 0.003 | -0.002 |
| 163 | *Laccophilus minutus* | 0.043 | -0.017 | 0.021 | 0.033 |
| 164 | *Laccophilus* sp. 01 | 0.027 | -0.041 | 0.023 | -0.006 |
| 165 | *Laccophilus* sp. 02 | -0.004 | -0.012 | 0.005 | 0.022 |
| 166 | *Laccophilus* sp. 03 | -0.008 | -0.012 | 0.005 | 0.017 |
| 167 | *Lestes viridis* | 0.053 | -0.089 | 0.053 | 0.007 |
| 168 | Libellulidae 01 | 0.001 | 0.008 | -0.003 | 0.009 |
| 169 | Limnephilidae 02 | -0.001 | 0.008 | -0.003 | 0.002 |
| 170 | *Limnoxenus niger* | -0.001 | 0.006 | 0.002 | -0.001 |
| 171 | *Micronecta* sp. 01 | 0.023 | -0.008 | 0.008 | 0.021 |
| 172 | *Microvelia* sp. 01 | 0.002 | -0.056 | 0.004 | -0.030 |
| 173 | *Microvelia* sp. 02 | -0.005 | -0.011 | 0.008 | -0.014 |
| 174 | *Musculium* sp.2 | -0.004 | -0.003 | -0.007 | 0.000 |
| 175 | *Musculium* sp.3 | 0.008 | 0.055 | 0.029 | 0.009 |
| 176 | *Myxas* sp. | -0.008 | -0.007 | 0.006 | -0.007 |
| 177 | *Naucoris maculatus* | -0.003 | 0.005 | 0.016 | -0.008 |
| 178 | *Noterus clavicornis* | 0.019 | -0.023 | 0.008 | 0.013 |
| 179 | *Notonecta* sp. 01 | -0.027 | 0.101 | 0.050 | 0.035 |
| 180 | *Notonecta* sp. 02 | 0.053 | -0.008 | 0.001 | 0.071 |
| 181 | *Ochthebius minutus* | -0.004 | -0.002 | -0.003 | -0.005 |
| 182 | *Odontomyia* sp.01 | -0.002 | -0.007 | 0.000 | -0.005 |
| 183 | *Odontomyia* sp.02 | -0.002 | -0.005 | 0.000 | -0.004 |
| 184 | *Oplodontha* sp. 01 | -0.003 | -0.003 | -0.004 | -0.001 |
| 185 | *Oplodontha* sp. 02 | -0.004 | -0.003 | 0.003 | -0.003 |
| 186 | *Orthetrum* sp. 01 | -0.003 | -0.010 | 0.001 | -0.007 |
| 187 | Orthoclaadiinae 01 | 0.055 | -0.008 | 0.013 | 0.059 |
| 188 | Orthoclaadiinae 02 | 0.006 | -0.020 | 0.043 | -0.002 |
| 189 | Orthoclaadiinae 03 | 0.056 | -0.094 | 0.127 | 0.091 |
| 190 | Orthoclaadiinae 04 | -0.052 | -0.076 | 0.046 | -0.015 |
| 191 | *Peltodytes caesus* | -0.006 | -0.020 | -0.004 | -0.010 |
| 192 | *Physella acuta* | -0.182 | -0.115 | -0.118 | -0.089 |
| 193 | *Pisidium* sp.1 | 0.040 | 0.042 | 0.060 | -0.020 |
| 194 | *Pisidium* sp.2 | 0.002 | 0.012 | -0.004 | 0.013 |
| 195 | *Planaria torva* | 0.004 | 0.030 | -0.024 | 0.070 |
| 196 | *Platycnemis* sp. 01 | -0.004 | 0.031 | 0.020 | 0.024 |
| 197 | *Plea minutissima* | 0.086 | -0.357 | 0.112 | -0.082 |
| 198 | *Podura aquatica* | -0.002 | 0.049 | 0.009 | -0.039 |
| 199 | Polycentropodidae 01 | -0.001 | -0.001 | -0.001 | 0.000 |
| 200 | *Potamopyrgus antipodarum* | -0.143 | -0.343 | -0.054 | -0.223 |
| 201 | *Proasellus* sp. | -0.128 | -0.114 | -0.227 | 0.052 |
| 202 | *Procambarus clarkii* | -0.002 | -0.003 | -0.002 | -0.001 |
| 203 | *Pyrrhosoma nymphula* | -0.015 | -0.036 | 0.012 | -0.057 |
| 204 | *Ranatra linearis* | -0.002 | 0.002 | 0.003 | -0.002 |
| 205 | *Rhantus suturalis* | -0.001 | 0.012 | -0.005 | 0.003 |
| 206 | Sciomyzidae 01 | 0.000 | 0.003 | 0.000 | -0.001 |
| 207 | *Segmentina nitida* | -0.002 | 0.020 | 0.000 | -0.006 |
| 208 | *Sialis* sp. 01 | -0.006 | 0.011 | -0.009 | 0.007 |
| 209 | Sigara 01 | -0.002 | -0.019 | -0.007 | -0.006 |
| 210 | Sigara 02 | -0.020 | 0.037 | -0.032 | 0.075 |
| 211 | Sigara 03 | -0.034 | -0.013 | -0.006 | 0.039 |
| 212 | Sigara 04 | -0.008 | -0.023 | 0.003 | -0.005 |
| 213 | Sigara 06 | -0.031 | -0.047 | -0.017 | -0.027 |
| 214 | *Sympetrum* sp.1 | -0.003 | 0.002 | -0.001 | -0.004 |
| 215 | *Sympetrum* sp.2 | -0.007 | -0.016 | 0.011 | -0.020 |
| 216 | *Sympetrum* sp.3 | 0.058 | -0.069 | 0.070 | -0.007 |
| 217 | Syrphidae 01 | -0.002 | -0.001 | -0.003 | 0.001 |
| 218 | Tabanidae 01 | 0.000 | -0.003 | 0.000 | -0.004 |
| 219 | Tabanidae 02 | -0.001 | -0.004 | 0.000 | -0.006 |
| 220 | Tanypodinae 01 | -0.029 | 0.120 | 0.031 | -0.039 |
| 221 | Tanypodinae 02 | -0.025 | 0.009 | 0.056 | -0.005 |
| 222 | Tanypodinae 03 | 0.000 | -0.037 | 0.013 | -0.020 |
| 223 | Tanypodinae 04 | -0.002 | -0.038 | 0.064 | -0.079 |
| 224 | Tanypodinae 05 | -0.002 | 0.047 | 0.031 | -0.042 |
| 225 | Tanypodinae 06 | -0.017 | 0.028 | 0.000 | -0.019 |
| 226 | Tanypodinae 07 | -0.001 | 0.021 | 0.008 | -0.015 |
| 227 | Tanypodinae 08 | -0.004 | -0.015 | 0.001 | -0.009 |
| 228 | Tanypodinae 09 | -0.018 | 0.022 | 0.000 | 0.000 |
| 229 | Tanypodinae 10 | 0.009 | -0.001 | 0.036 | -0.035 |
| 230 | Tanypodinae 11 | -0.004 | 0.039 | 0.014 | -0.001 |
| 231 | *Tanysphyrus lemnae* | -0.001 | 0.014 | 0.009 | -0.013 |
| 232 | Tanytarsini 01 | -0.053 | 0.014 | 0.078 | 0.049 |
| 233 | Tanytarsini 02 | -0.007 | 0.004 | 0.006 | -0.010 |
| 234 | *Theremyzon* sp. 01 | -0.006 | -0.002 | 0.000 | -0.008 |
| 235 | Trichoptera 02 | -0.004 | -0.002 | -0.006 | 0.001 |
| 236 | *Valvata macrostoma* | 0.495 | 0.153 | 0.105 | 0.459 |

